# Cleave and Rescue gamete killers create conditions for gene drive in plants

**DOI:** 10.1101/2023.10.13.562303

**Authors:** Georg Oberhofer, Michelle L. Johnson, Tobin Ivy, Igor Antoshechkin, Bruce A. Hay

## Abstract

Gene drive elements promote the spread of linked traits, even when their presence confers a fitness cost to carriers, and can be used to change the composition or fate of wild populations. Cleave and Rescue (*ClvR*) drive elements sit at a fixed chromosomal position and include a DNA sequence-modifying enzyme such as Cas9/gRNAs (the Cleaver/Toxin) that disrupts endogenous versions of an essential gene, and a recoded version of the essential gene resistant to cleavage (the Rescue/Antidote). *ClvR* spreads by creating conditions in which those lacking *ClvR* die because they lack functional versions of the essential gene. We demonstrate the essential features of *ClvR* gene drive in the plant *Arabidopsis thaliana* through killing of gametes that fail to inherit a *ClvR* that targets the essential gene YKT61, whose expression is required in male and female gametes for their survival. Resistant (uncleavable but functional) alleles, which can slow or prevent drive, were not observed. Modeling shows plant *ClvR*s are likely to be robust to certain failure modes and can be used to rapidly drive population modification or suppression. Possible applications in plant breeding, weed control, and conservation are discussed.

## Main

Gene drive occurs when genetic elements—genes, gene complexes, large chromosomal regions or entire chromosomes—are transmitted to viable, fertile progeny at rates greater than those of competing allelic variants or other parts of the genome. There has long been interest in the idea that genetic manipulation of wild populations via gene drive could be used for beneficial purposes. Transgenes or alleles of endogenous loci can be linked with a drive element. The results of modeling and lab experiments demonstrate this can result in spread of these “cargo” genes to high frequency in an extant population. Alternatively, drive can result in population suppression or elimination if spread of the element drives the population towards an unfit set of genotypes (e.g., all male, all females sterile), reviewed in^1–3^.

A number of applications of gene drive in plants have been proposed^4–6^. In the context of agriculture, gene drive has been discussed as a way to spread desirable agronomic traits, and as a possible tool for weed management, by sensitizing the population to some other form of intervention or by suppressing it directly. Ecosystem engineering/conservation is another possibility. This could take the form of suppressing invasive species. Alternatively, population modification could be used to engineer pathogen resistance or other forms of resilience (a form of evolutionary rescue) into native species in the face of current stresses—or anticipated novel stresses due to climate change.

A variety of selfish genetic elements have been considered for bringing about gene drive. These include transposons and homing endonucleases, which spread through over-replication; multigene complexes that produce female meiotic drive or sperm post-meiotic segregation distortion; and toxin-antidote combinations that spread by causing the death of those (cells, spores, gametes, or progeny) who fail to inherit them from a carrier. Toxin-antidote gene drive elements (TA elements) are particularly interesting as they are found throughout all domains of life: prokaryotes, fungi, animals and plants and the wide distribution of some of these elements in nature shows they can spread and persist in complex natural environments^7–9^. Here we focus on eukaryotes and drive associated with sexual reproduction.

A TA element sits at a fixed chromosomal position and consists of one or more genes that encode linked toxin and antidote functions. The toxin, typically a protein, has the potential to kill or impair the development of those in which it is present, while the antidote, a protein or RNA, suppresses the activity or expression of the toxin^7,10–12^. The toxin is trans-acting and is distributed to all meiotic products or progeny of a TA-bearing parent. However, only those that inherit the TA cassette express the antidote, which counteracts the toxin in cis. In consequence, TA elements ensure their presence in the next generation by causing the death of those who fail to inherit them (post-segregational killing) from a parent, a form of genetic addiction. The death of those lacking the TA cassette can result in a relative increase in frequency of those carrying it. Modeling shows that TA elements in sexually reproducing eukaryotes can (depending on the fitness costs associated with carriage of the element and introduction frequency) spread to high frequency even if they do not confer any advantage to their hosts^13–23^.

TA elements in nature^7,10–12,24^, including those in plants ^9,12,25–33^ evolved in specific genomic, organismal and ecological contexts, and it is often unclear if the mechanisms of action, associated gene regulation and species-specific information on development (timing and levels of gene and protein expression and localization) can be easily transferred to bring about drive in other species. Similar considerations apply to synthetic *Medea* TA elements engineered in *Drosophila* in which the toxin is an engineered transient loss-of-function (LOF) of a maternally expressed gene whose product is essential for embryogenesis and the antidote is a zygotically expressed transgene that restores this missing function in a just-in-time fashion^34–36^.

Recently, in an effort to create a chromosomal TA-based gene drive system that utilizes a LOF toxin and consists of a simple and extensible set of components that can plausibly be implemented across diverse species we developed the *Cleave and Rescue* (*ClvR*) element^19,37–39^, also referred to as Toxin Antidote Recessive Embryo (TARE)^20,40^ in related implementations (hereafter referred to as *ClvR*, a name that captures the key mechanisms involved). A *ClvR* element encodes two activities. The first component, the Cleaver/Toxin, is a DNA sequence–modifying enzyme such as Cas9 and multiple guide RNAs (gRNAs). These are expressed in the germline or cells that will become the germline, though germline-specific expression is not required. Cas9 and its associated gRNAs disrupt—through cycles of cleavage and end joining that continue until the target site is destroyed—endogenous versions of a haplosufficient (and in some contexts haploinsufficient or haplolethal) essential gene, wherever it is located. Inaccurate repair at multiple positions in the coding region of the essential gene creates loss-of-function (LOF) alleles. These are the potential toxin. The second component of *ClvR*, the Rescue/Antidote, is simply a recoded version of the essential gene resistant to cleavage and gene conversion with the cleaved version, expressed under the control of regulatory sequences sufficient to rescue the LOF phenotype. LOF alleles of the essential gene, which segregate and exist independently of *ClvR*, perform their toxin function when they find themselves (potentially many generations later) in homozygotes that die because they lack the *ClvR*-derived source of essential gene function. In contrast, those who inherit *ClvR* and its associated Rescue survive. In this way, as with TA-based selfish genetic elements found in nature, *ClvR* increases in relative frequency by causing the death of those that lack it. This results in cells, organisms, and ultimately populations becoming dependent on (addicted to) the *ClvR*-encoded Rescue transgene for their survival.

In *Drosophila*, autonomous *ClvR*/TARE elements have been created and shown to spread in wildtype (WT) populations to transgene fixation (all individuals carry at least one copy)^19,20,37,39,40^. Other features, such as the ability to create strong, but self-limited drive^38^, engage in multiple cycles of population modification that replace old content with new^37^, and achieve population suppression using a conditional Rescue^39^ have also been demonstrated. Multiple other configurations of the components that make up *ClvR*/TARE have been proposed, and modeling predicts they can give rise to drive with a diversity of interesting characteristics for population modification or suppression^21,41^.

In animals *ClvR*-type drive is most easily implemented through the killing and rescue of specific zygote genotypes, as above in *Drosophila*. Engineering TA drive based on killing and rescue of gametes is also of great interest because gametic drive can be much stronger than zygotic drive. It will typically drive the element to allele fixation (as opposed to transgene fixation with many zygotic TA elements, which includes heterozygotes), and it can be used to bias sex ratios if the driver is linked to a sex chromosome and has its effects during post-meiotic spermatogenesis. These latter two features are important for several proposed methods of population suppression ^13,21,42^. A number of naturally occurring gametic drive systems (most often biasing sperm genotypes) in animals have been characterized, but the information available does not yet provide guidance as to if or how they can be used as tools ^43,44^. Engineering *ClvR*-based gametic drive in animals is challenging for several reasons. In females the gamete is chosen through differential segregation of one of the products of meiosis within the common cytoplasm of the oocyte and thus there is no opportunity for the antidote to select for carriers. In spermatogenesis the haploid spermatid products of a meiosis are connected by cytoplasmic bridges until late in development, and active content sharing of many but not all products (e.g.^45^) limits opportunities for bringing about differential killing and survival^46^.

The same strategies used to build *ClvR* in animals (killing of specific zygote genotypes) could also be implemented in plants. In addition, in our original description of *ClvR* we noted that gametic drive could be implemented in sexual organisms such as fungi and plants (Fig. S1 in ^19^, and ^47^), in which sibling gametes do not share components and require haploid gene expression for development and/or survival. In plants, meiotic products undergo additional rounds of mitosis, developing into multicellular haploid gametophytes (the female megagametophyte and male microgametophyte) that produce ovules or pollen. This requires extensive expression of the haploid genome^48^. These features of plant gamete development are reflected in the many recessive mutants (no somatic phenotype in the heterozygous diploid or polyploid parent; the sporophyte stage) that cannot be transmitted through one or the other sex, often identified through sex-specific transmission ratio distortion (e.g.^49,50^). Mutations in other genes cannot be transmitted through either sex due to a requirement in both gametophyte types^51–53^. These characteristics make plants an ideal system in which to implement gene drive based on a *Cleave and Rescue* mechanism in which gametes die if they fail to inherit *ClvR* from a *ClvR*-bearing parent. (Fig. 1a-d).

**Fig. 1:**
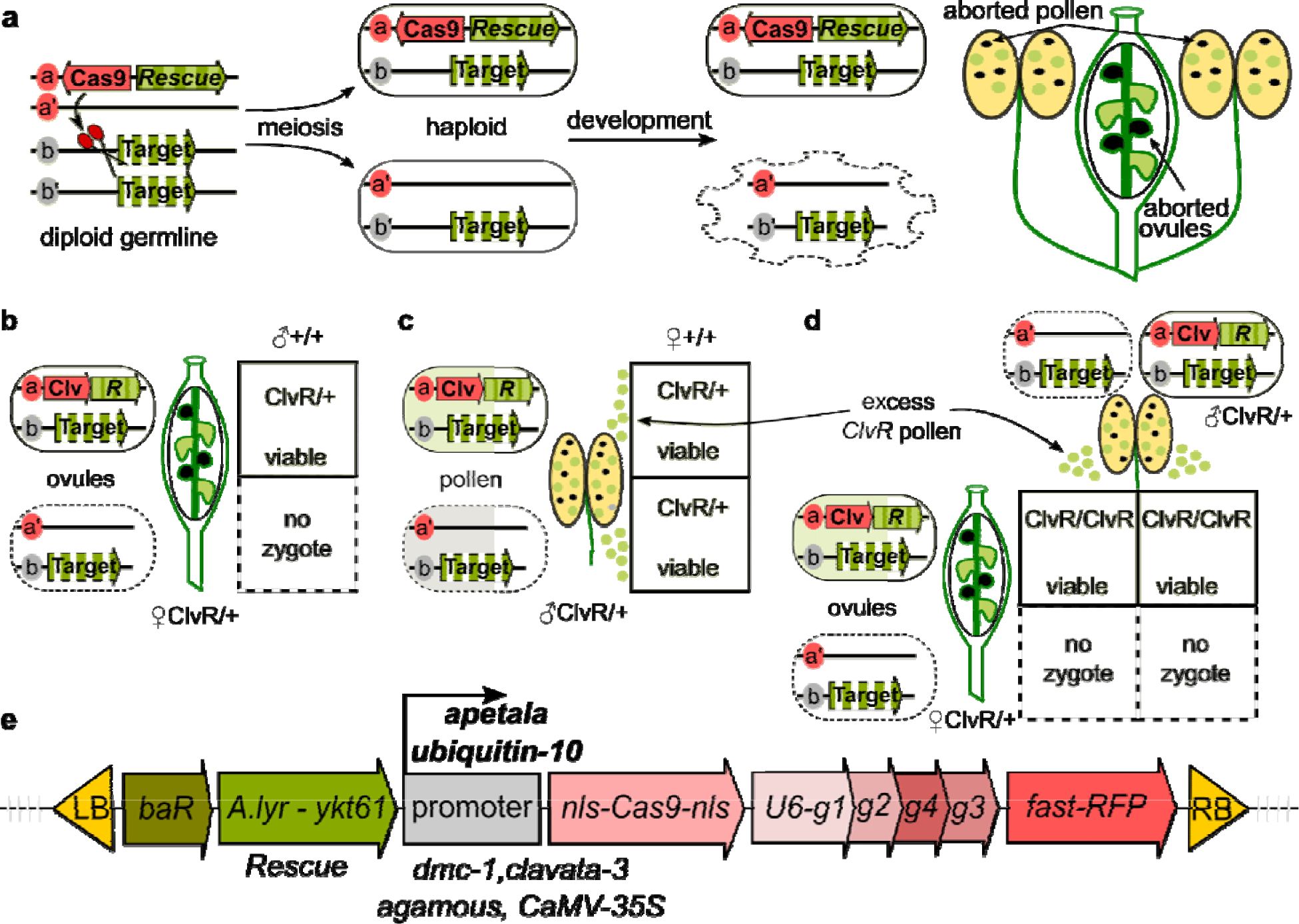
*ClvR* behavior in a diploid plant and construct design. **(a)** Cas9/gRNAs located on chromosome **a** of an **a/a’** *ClvR* heterozygote cleave an essential gene located on chromosome **b** and **b’** during the diploid parental stage, creating LOF alleles. Diploid cells survive this because the *ClvR* carries a recoded rescuing version of the essential gene, which produces a functional product (light green background). During haploid stages expression of the essential gene is required for gamete/gametophyte development/survival. Gametes/gametophytes that fail to inherit *ClvR* lack a functional copy of the essential gene and die (indicated with dashed outline and gray background). The hermaphrodite plant to the right has anthers with *ClvR*-bearing pollen (green circles) and dead non-*ClvR*-bearing pollen (dark circles) and an ovary containing *ClvR-*bearing ovules (large green shapes) and dead non-*ClvR*-bearing ovules (large dark circles). **(b)** Cross of a heterozygous *ClvR*-bearing female with WT (+/+) pollen. Non-*ClvR-*bearing gametophytes die and do not undergo fertilization (gray no zygote square). Thus, all progeny are *ClvR*-bearing heterozygotes (green square). **(c)** Cross in which pollen from a heterozygous *ClvR*-bearing male fertilizes ovules of a WT (+/+) female. Pollen is produced in large excess over ovules. Thus, death of the 50% non-*ClvR*-bearing pollen (dark circles) still allows all ovules to be fertilized, resulting in all progeny being *ClvR*-bearing heterozygotes. **(d)** Cross of a heterozygous *ClvR* female to a heterozygous *ClvR* male. Only *ClvR*-bearing ovules and pollen participate in fertilization, resulting in all progeny being homozygous *ClvR/ClvR*. **(e)** Genetic makeup of the *ClvR* drive element. From left to right these are: a Basta herbicide resistance marker (baR); a YKT61 rescue transgene derived from *Arabidopsis lyrata* (*A. lyrata* - YKT61); one of six different enhancer/promoters (those that resulted in significant transmission ratio distortion indicated in bold) used to direct Cas9 expression; Cas9 (one of two different versions, discussed in text); 4 gRNAs designed to base pair with DNA for the YKT61 coding region, with each expressed under the control of an independent U6 promoter (U6-g1-4); a fluorescent seed transgenesis marker (fast-RFP). Repeats required for transgenesis using agrobacterium (LB and RB) flank these elements.

*Arabidopsis thaliana* is a good system in which to test self-sustaining gene drive constructs in plants because it is a self-fertilizing hermaphrodite in which fertilization typically occurs before flower opening, thus limiting opportunities for pollen/gene flow. In addition, *A. thaliana* is not naturally wind pollinated and lab and field experiments demonstrate that outcrossing rates are very low^54–58^. Thus, transgene containment is straightforward. However, for these same reasons population level gene drive experiments of the type carried out in insects—mixed populations of transgenic and non-transgenics allowed to mate freely and followed for changes in genotype frequency over multiple generations—cannot be carried out. Here we show, using manual mating between parents of different genotypes, the key features required for *ClvR* drive: a high frequency of LOF allele creation, a high, non-Mendelian rate of *ClvR* inheritance in progeny, and the absence of resistant alleles (mutated, uncleavable, but functional) that would slow or subvert the intended goal of drive, population modification or suppression. Modeling shows that elements with the features we demonstrate experimentally have the potential, in diploid obligate outcrossing species (dioecious), to bring about population modification or suppression. Possibilities for drive in hermaphrodites (male and female reproductive organs in the same flower) and monoecious species (male and female flowers on the same plant), and the consequences of inbreeding, are also discussed. Together our observations, along with those of Liu and colleagues in related work^59^, suggest possible applications, as well as challenges, for use of *ClvR* gamete killer gene drive in plants.

## Results

### Components of a *ClvR*-based gamete killer

The strength of a gene drive—its ability to spread from low frequency and in the presence of significant fitness costs—is increased when it biases inheritance in its favor in both sexes, something that is of particular importance when trying to bring about population suppression. Engineering *ClvR*-based gamete drive with this feature (Fig. 1a) requires targeting a gene whose expression during the haploid stage is required for the survival and/or development of the microgametophyte (referred to as pollen, which contains sperm) and megagametophyte (referred to as ovule, a sporophytic structure in the ovary within which each megagametophyte, which includes the egg, develops). Mutations in many ubiquitously expressed housekeeping genes (such as were targeted for LOF allele creation in insect *ClvR*s^19,37,40^), likely have such a phenotype in plants given the extensive gene expression that occurs in gametes, but are challenging to identify since such mutations cannot be passed through the germline. Their identity is sometimes inferred by their absence in mutant collections (e.g.^51^). Alternatively, with the advent of methods for CRISPR-based mutagenesis, genes whose mutation results in loss of male and female gametes can be identified through reverse genetics approaches that incorporate a rescuing transgene into the mutagenized genetic background (e.g.^52,53^). Here we focus on one such gene, YKT61, a ubiquitously expressed R-SNARE protein involved in fusion between vesicle and target membranes^52^. Formally, it is not known if the YKT61 LOF phenotype in sporophytes is recessive lethal since crosses between heterozygotes do not produce viable LOF gametes^52^. That said, YKT61 is expressed ubiquitously and recent work shows that partial loss of function of YKT61 using RNA interference has strong effects on root development^60^. Thus, its loss of function is likely to be at least deleterious in the sporophyte.

The components that make up our *ClvR* gamete killers are illustrated in Fig. 1e. As a Rescue we utilized a genomic fragment containing the *Arabidopsis lyrata* YKT61 gene (in which some amino acid coding region differences were recoded back to those of *A. thaliana*, Extended Data Fig. 1). For the Cleaver, four gRNAs targeting conserved regions within the *A. thaliana* YKT61 coding sequence (see also Fig. 4) were expressed ubiquitously using individual U6 Pol-III promoters^61^. Several versions of Cas9 were tested. One lacks introns and carries a mutation (K918N) shown to increase Cas9 catalytic activity^62^ while a second one contains 13 introns, which are thought to increase expression^63^. Regulatory sequences from six different genes were used to direct Cas9 expression. *Arabidopsis* DMC1 is primarily expressed during meiotic stages^64^. Sequences from the CLAVATA3 (early stem cell identity^65^), APETALA1 (flower meristem identity^66^) and AGAMOUS (reproductive floral organ primordia^67^) genes direct expression in adult sporophyte tissues that include the future germline. The CaMV35S^68^ and UBIQUITIN10^69^ promoters direct expression broadly, in many if not all cell types. The DMC1 promoter was used in combination with both versions of Cas9, while AGAMOUS, CLAVATA3, APETALA1, CaMV35S and UBIQUITIN10 sequences were used to direct expression of the version of Cas9 lacking introns.

### *ClvR*s based on cleavage and rescue of the YKT61 gene show key features required for gamete killer gene drive

We used floral dipping with agrobacterium to transform a number of T0 WT plants with the above constructs (Fig. 2a). A number of independent transformants, identified as red transgenic seeds of the T1 generation, were collected from these plants (Fig. 2b) and characterized in the crosses outlined in Fig. 2c-f. T1 seeds (heterozygous for one or more *ClvR* elements) were grown to adulthood and allowed to self (T1xT1; Fig. 2b). T1 self crosses that produced progeny siliques (a seed pod, which contains progeny from the ovules of one flower) containing all or primarily red seeds—the T2 generation, possibly *ClvR/ClvR* homozygotes (Fig. 2c)—were characterized further as this is the expected phenotype if gametic drive occurred in one or both sexes in a cross between heterozygotes (Fig. 1b-d). Based on the results of these experiments (a significant fraction of non-*ClvR* seeds), constructs utilizing regulatory sequences from the DMC1, AGAMOUS, and CLAVATA-3 genes were not considered further.

**Fig. 2.**
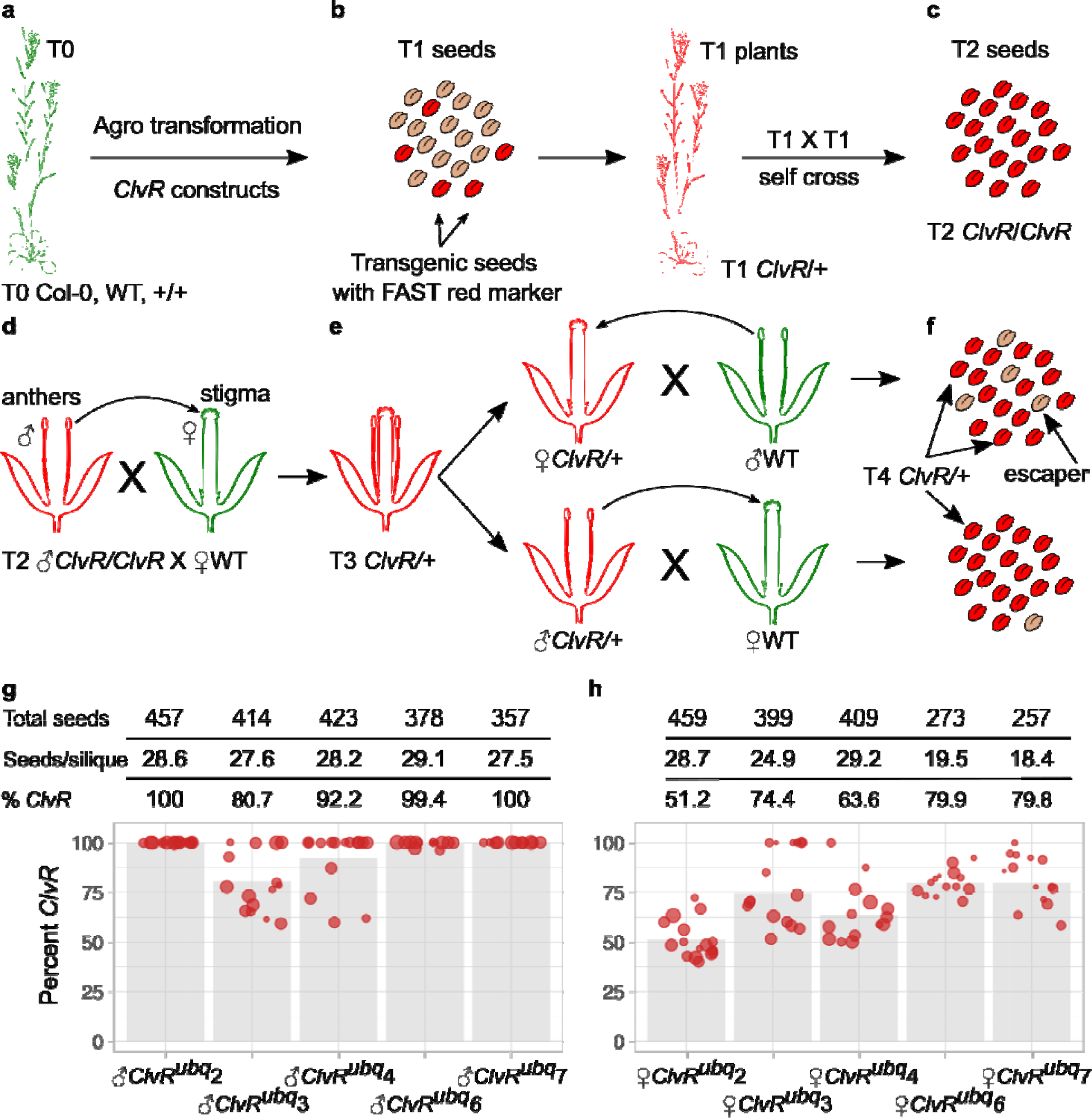
Genetic evidence for *ClvR*-based gamete killing and rescue. **(a-f)** Crosses used to establish independent *ClvR* insertions **(a-d)** and test for gametic drive through the male and female germline **(e-f)**. **(g,h)** Frequency of *ClvR* transmission through the male **(g)** and female **(h)** germline. We counted the total number of seeds and the number of seeds per silique. Each circle represents an individual silique. The size of the red circle scales with the number of seeds in the silique. Gray bars represent mean *ClvR* inheritance values from all seeds in a cross. Note that for the female crosses in **h**, in general as the frequency of *ClvR* inheritance goes up, the number of seeds in the silique goes down. This is expected since the number of functional ovules determines the maximal seed output, and a *ClvR* with efficient killing and rescue would only be expected to produce half the WT number of functional ovules/seeds. Seed and silique counts are in Supplementary Table S1.

T2 seeds carrying constructs that utilized the APETELA1 and UBIQUITIN10 regulatory sequences were grown to adulthood and pollen from these plants was used in an outcross to WT to produce T3 *ClvR*/+ seeds (Fig. 2d). Finally, in the key outcross to test for gametic drive, T3 seeds were grown to adulthood and pollen and ovules used in outcrosses to WT (Fig. 2e). The frequency of *ClvR* inheritance (*ClvR/*+) in progeny T4 seeds provides a measure of gamete killing and rescue (Fig. 2f). *ClvR* inheritance rates in T4 seeds from T3 pollen, shown for 5 different insertions using UBIQUITIN10 sequences (*ClvR*^ubq^ lines) to direct Cas9 expression (Fig. 2g) were generally very high, with three of the five showing inheritance rates greater than 99%. Inheritance rates of *ClvR*^ubq^ in T4 seeds from T3 ovules were also significantly above 50%, but a number of non-*ClvR* seeds (generically referred to as escapers) were observed (Fig. 2h). Similar results were obtained for *ClvR*s utilizing APETALA1 regulatory sequences (Extended Data Fig. 2). Crosses with *ClvR*s using the CaMV35S promoter showed inheritance that was modestly *ClvR*-biased (Extended Data Fig. 3). These were not considered further. The basis for the differences between drive through pollen and ovules is addressed further below.

### *ClvR*-based gamete killing and rescue is stable over multiple generations

The above results show that *ClvR*s designed to kill and rescue a gamete essential gene can bias inheritance in their favor in *Arabidopsis*, satisfying the key requirement for TA-based gametic gene drive. We focused our characterization on the *ClvR*^ubq7^ line as it is associated with a single insertion, showed a high frequency of inheritance through pollen and ovules, and heterozygotes and homozygotes were otherwise healthy (Extended Data Fig. 4). To determine if the bias in *ClvR*^ubq7^ inheritance was stable, and if it had any dependence on the sex through which drive element was inherited, we characterized drive of *ClvR*^ubq7^ alleles present in T4 *ClvR*^ubq7^/+ individuals (Fig. 2f) that came from either a male or female *ClvR*^ubq7^/+ T3 parent (Fig. 3). *ClvR*^ub*q*7^-bearing T4 seeds derived from male or female *ClvR*^ubq7^*/*+ T3 parents crossed to WT were grown to adulthood and pollen and ovules from *ClvR*^ubq7^-bearing T4 individuals were used in outcrosses to WT, giving rise to a T5 generation of seeds whose *ClvR*^ubq7^/+ grandparents (the T3 generation) and parents (the T4 generation) were either both female (ovules), both male (pollen), or one (T3) and then the other (T4) (Fig. 3a,b). As shown in Fig. 3c,d inheritance rates remained comparable – very high when transmitted through pollen and high but with significantly more non-*ClvR*^ubq7^-bearing escaper seeds when transmitted through ovules – regardless of the parental and grandparental sex.

**Fig. 3.**
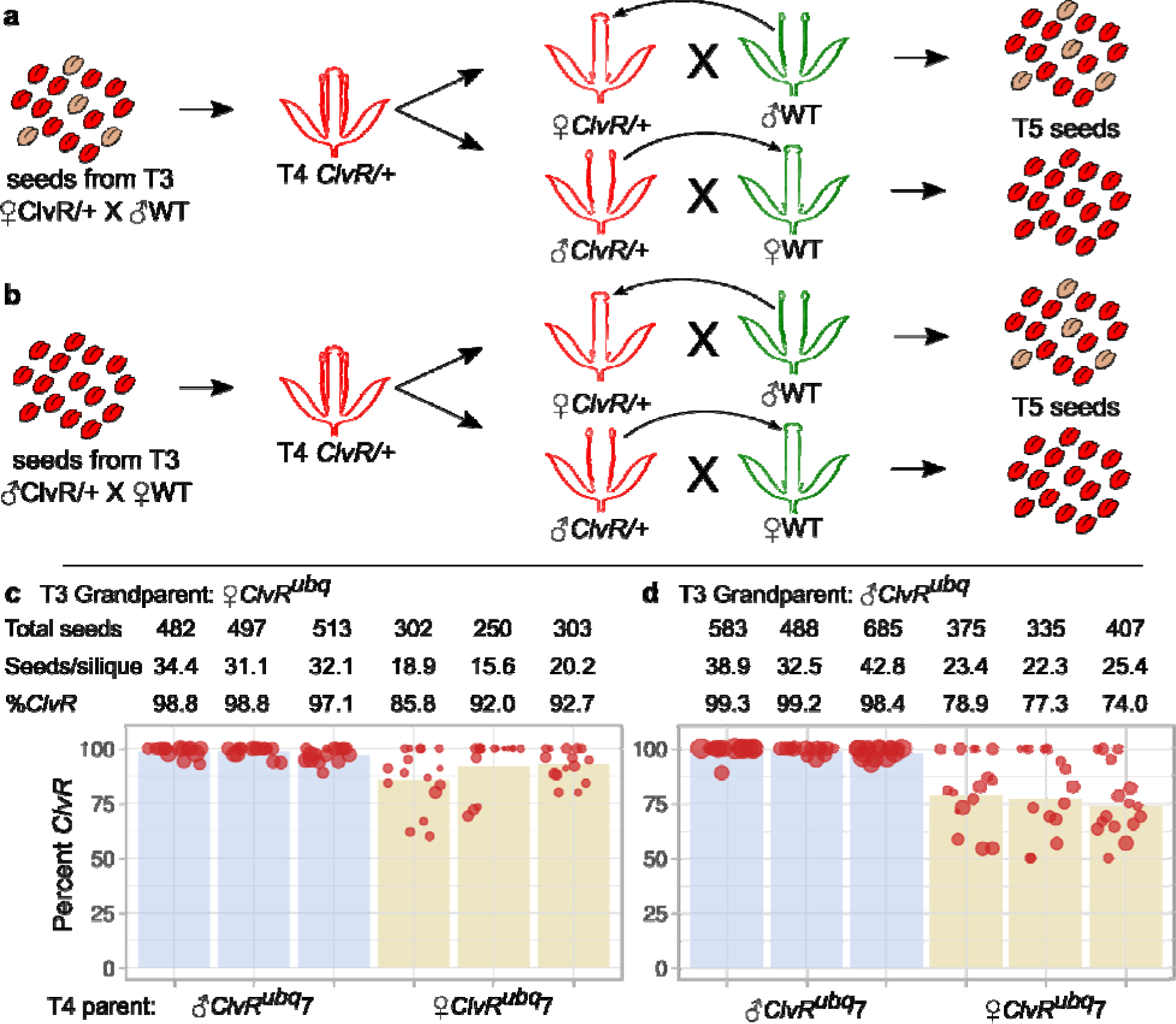
*ClvR*-based gamete killing and rescue is stable over multiple generations. **(a-b). (a)** A T3 cross between *ClvR*^ubq7^/+ females and WT was used to generate T4 *ClvR*^ubq7^/+ heterozygous plants. Pollen and ovules from these T4 individuals were used in outcrosses to WT to generate a *ClvR*^ubq7^ heterozygous T5 generation. **(b)** A T3 cross between *ClvR*^ubq7^*/*+ males and WT was used to generate T4 *ClvR*^ubq7^/+ heterozygous plants. Pollen and ovules from these T4 individuals were used in outcrosses to WT to generate a *ClvR*^ubq7^ heterozygous T5 generation. **(c)** Frequency of *ClvR*^ubq7^ inheritance in crosses in which the T3 grandparent was a *ClvR*^ubq7^/+ heterozygote female and T4 parents were either a *ClvR*^ubq7^/+ female (left six columns) or male (right six columns). **(d)** Frequency of *ClvR*^ubq7^ inheritance in crosses in which the T3 grandparent was a *ClvR*^ubq7^/+ heterozygote male and T4 parents were either female (left six columns) or male (right six columns). Seed and silique counts are in Supplementary Table S1. A description of crosses is presented in Extended Data Fig. 5.

### Mutations associated with cleavage and molecular basis of escape from gamete killing

The high frequency of *ClvR*^ubq7^ inheritance when transmitted through *ClvR*^ubq7^/+ pollen argues that rates of cleavage and LOF mutation creation are high, and that rescue is efficient. The UBIQUITIN10 regulatory sequences drive expression broadly throughout development, from the embryo onwards^70^, long before the male and female germlines form. This suggests that rates of cleavage and LOF allele creation in female gametes are high as well. To understand the molecular events associated with drive, and the unexpectedly high numbers of non-*ClvR*^ubq7^ progeny observed when a *ClvR*^ubq7^*/*+ individual was the female parent, we sequenced the endogenous YKT61 locus in leaves of several genotypes: *ClvR*^ubq7^/+ T4 heterozygotes and *ClvR*^ubq7^/*ClvR*^ubq7^ T5 homozygotes derived from a T4 self cross; and non-*ClvR*^ubq7^ escapers from crosses of *ClvR*^ubq7^/+ to WT, in which the *ClvR*^ubq7^/+ was the female or male parent (Fig. 4). See Extended Data Table 1 for the details of sequence alterations at each gRNA target site. In *ClvR*^ubq7^/+ individuals all four target sites were cleavable and mutated to LOF (frameshifts) at high frequency, with at least four sites being altered in all five plants. In *ClvR*^ubq7^*/ClvR*^ubq7^ homozygotes all four sites were altered in all four sequenced individuals. In outcrosses using *ClvR*^ubq7^/+ pollen a very small number of non-*ClvR*^ubq7^ escaper seeds was observed (∼1% of all seeds; Fig. 2 and Fig. 3). Six T5 escapers were grown to adulthood and sequenced. All were WT at all four gRNA target sites (Fig. 4). Thus, escape from death by non*-ClvR*^ubq7^ pollen is due to lack of cleavage and/or sequence alteration following cleavage, at all four sites. In contrast, in crosses with females as the parent nine out of ten non-*ClvR* progeny of a *ClvR*^ubq7^/+ parent carried one or more sequence alterations at gRNA target sites that create LOF alleles (frameshifts) in the YKT61 coding region (Fig. 4 and Extended Data Table 1). The frequency of mutation at each target site in escapers from a female *ClvR* parent is reduced as compared with that observed in *ClvR/*+ and *ClvR/ClvR* genotypes. However, this is is expected since mutagenesis is ongoing in the *ClvR* carriers but not escapers. Finally, the other escaper was wildtype at all four target sites. No resistant versions of endogenous YKT61—mutated but likely to be functional—were observed.

**Fig. 4:**
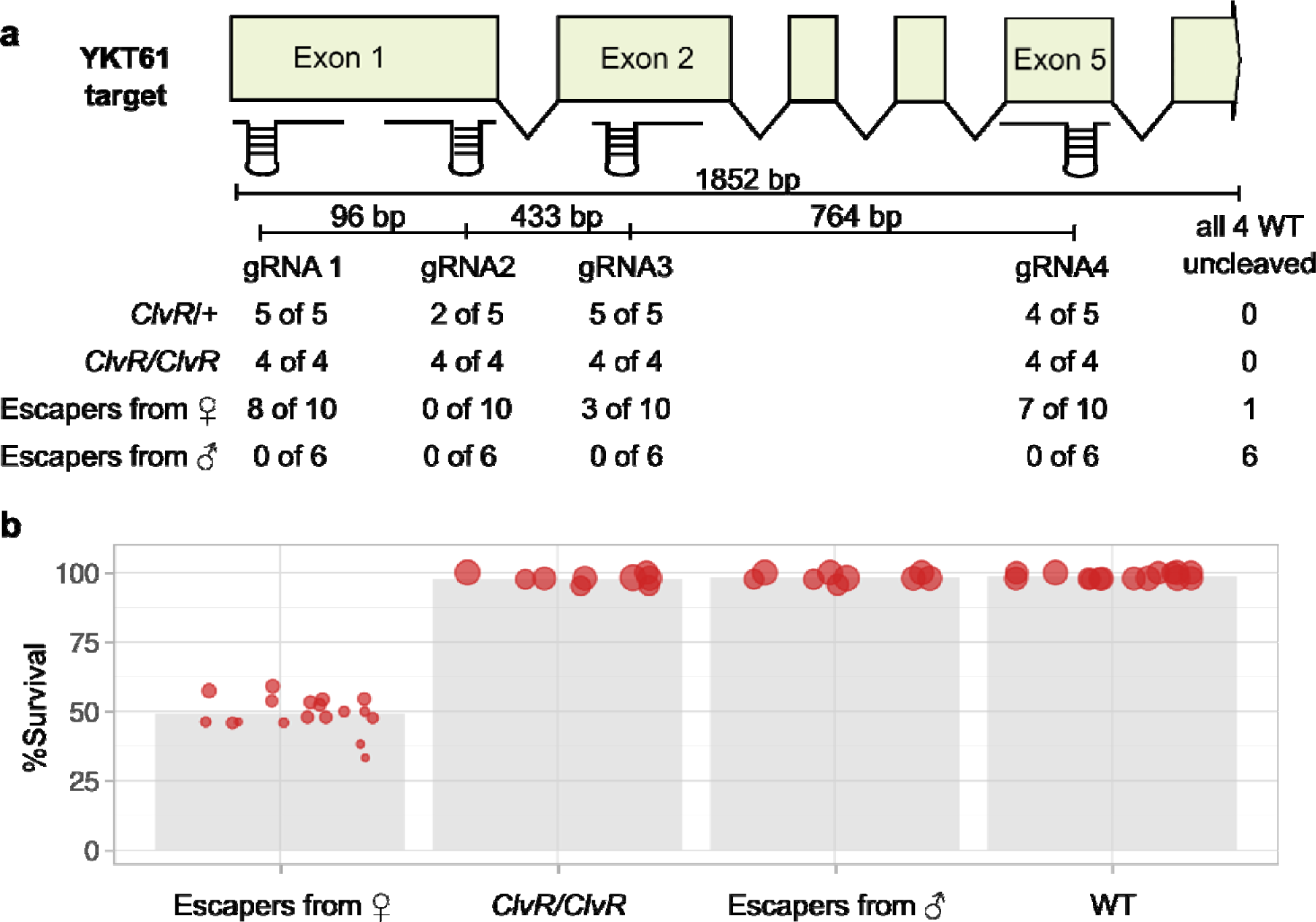
Characterization of the target locus following exposure to *ClvR*^ubq^^7^, and genetic behavior of LOF mutations found in female escapers. **(a)** The genomic region containing the YKT61 gene is shown, along with the approximate locations of four gRNA target sites (see Extended Data Fig. 1 for exact sequences). The table summarizes how many of the target sites were altered to likely LOF (frameshifts or large deletions) in *ClvR*^ubq7^/+, *ClvR*^ubq7^*/ClvR*^ubq7^ (heterozygous *ClvR*^ubq7^/+, inbred for one generation), escapers coming from a *ClvR*^ubq7^/+-bearing mother, and escapers from a *ClvR*^ubq7^-bearing father. One out of ten escapers from *ClvR*^ubq7^ mothers were WT at all four target sites. Six out of six escapers from *ClvR*^ubq7^ fathers were WT at all four target sites. **(b)** Fraction of total ovules in individual siliques (red circles) that developed into seeds in self crosses from 4 different genotypes. Size of the circle scales with number of seeds in a silique. Escapers derived from a female *ClvR*^ubq7^/+ parent aborted ∼50% of ovules, consistent with the known LOF phenotype of YKT61 mutants^52^. Escapers derived from a male *ClvR*^ubq7^ parent (all WT at the YKT61 locus) showed high seed production as did *ClvR*^ubq7^/*ClvR*^ubq7^ homozygotes.

To summarize, rates of cleavage and mutation to LOF at the YKT61 locus are very high in male and female gametes. In males death of non-*ClvR*^ubq7^ pollen, coupled with efficient rescue of those inheriting *ClvR*^ubq7^, leads to a very high frequency of *ClvR*^ubq7^ inheritance in progeny. In females inheritance of *ClvR*^ubq7^ is also high, but there are also significant numbers of non-*ClvR*^ubq7^ progeny. The results of sequencing show that most of these have a LOF mutation, probably created by *ClvR*^ubq7^ at a much earlier stage in the diploid sporophyte. The construct was not present in plants grown from escaper seeds (Extended Data Fig. 6), arguing against transgene silencing playing a role. In earlier work, Cas9-induced LOF alleles of YKT61 were uniformly not passed to progeny through female gametes, resulting in abortion of 50% of ovules in a self cross^52^. Given this, our observations suggest that in the *ClvR*^ubq7^/+ diploid the YKT61 rescue transgene from *A. lyrata* provides YKT61 transcript and/or protein that is carried over from the mother into the non-*ClvR*^ubq7^ haploid ovules (maternal carryover rescue), and that this is sufficient to rescue the survival of some gametes carrying a LOF YKT61 allele. A strong prediction of this hypothesis is that LOF alleles generated by *ClvR*^ubq7^ and present in LOF/+, non-*ClvR*^ubq7^ heterozygotes should, since they lack the *ClvR^ubq^*^7^ Rescue, not be transmitted to the next generation. In a self cross of female escapers (most of whom are LOF/+ heterozygotes based on the results of sequencing; Fig. 4) this should manifest itself as 50% abortion in progeny siliques. As illustrated in Fig. 4b and Extended Data Table 2 this is the phenotype we observed for a number of female escapers tested. In contrast, and as expected, the ovule abortion rate in self crosses of homozygous *ClvR*^ubq7^/*ClvR*^ubq7^, male escapers, and WT was very low.

The mechanism by which maternal *ClvR* rescues some female gametes from death requires further exploration. Recoding associated with use of *A. lyrata* YKT61 may have created an mRNA with an extended half-life. Alternatively, position effects based on chromatin structure and/or nearby transcriptional regulatory sequences may lead to increased expression and/or extend expression of *A. lyrata* YKT61 farther into meiosis, resulting in carryover into non-*ClvR* bearing gametes carrying a LOF mutation in YKT61. Incorporation of chromatin insulators^71^ and/or targeting of other essential genes can minimize such effects.

### Modeling suggests conditions under which *ClvR* elements can drive population modification and suppression in diploid plants

Plants, being immobile, have limited control over who they mate with, a process mediated by wind, water or pollinators. Gene drive can only occur in the presence of outcrossing, which provides an opportunity for different alleles to compete for transmission to viable and fertile progeny. Thus, the potential for gene drive is minimal in self-fertilizing hermaphrodites and maximal in those that engage in obligate outcrossing. Seed-bearing plants (gymnosperms and angiosperms) have a variety of mating systems^72^, which can also vary among populations^73^. Most flowering plants (angiosperms; ∼95%) produce male and female gametophytes on the same plant. Many of these are hermaphrodites, with male and female gametophytes in the same flower (perfect flowers), while others are monoecious, with male and female flowers (imperfect flowers) on the same plant. For both these systems (in some but not all species), anatomical features (herkogamy), differences in the time of maturation of male and female gametophytes (dichogamy), or genetic forms of incompatibility, can reduce the likelihood of inbreeding^74,75^. Finally, a modest number of species (both gymnosperms and angiosperms) have separate sexes (dioecy), with male and female flowers on separate plants. Here we focus our modeling on dioecious species, in which outcrossing is obligate. Similar principles will apply to hermaphrodites and monoecious species, though inbreeding will always work to slow or prevent drive by reducing the frequency of outcrossing, which allows ClvR-associated fitness costs to accumulate and prevents them from being counterbalanced by the relative fitness increase gained from killing of non-*ClvR*-bearing alleles. These last points notwithstanding, it is noteworthy that a number of protein-based TA elements have been identified in rice^9,12,25,26,28–33^ a monoecious species with a high inbreeding coefficient^76^, and that one of these, the DUYAO-JIEYAO element, has spread to high frequency over the last 50 years^9^.

To explore the utility of *ClvR*-based gamete killers for population modification and suppression we used a stochastic model (a dioecious panmictic population with non-overlapping generations that considers individual and gametes; see methods for details) to explore *ClvR* behavior in several scenarios. This type of model is often used to gain insight into population genetic processes and provides a format that allows comparison of gene drive methods with respect to their basic population genetic features. However, it provides only heuristic guidance and is not predictive for any particular species or environment since it does not include consideration of many other environment- and species-specific variables. These include the mating system, level of inbreeding, overlapping generations (including the presence of seed banks), spatial structure, pollen and seed flow throughout that structure, whether pollen is in excess or limiting for fertilization, and the details of density dependence. Temperature sensitivity of DNA sequence modifying enzymes such as Cas9 and how this interfaces with climate and the timing of gamete development and cleavage will also be important to consider.

In animals, gene drive is often modeled using a paradigm in which matings are monogamous and male gametes (sperm) is not limiting. However, in many plants of interest (crops, weeds, targets of conservation) polyandry (fertilization of a female with pollen sourced from multiple males, which is in excess) and polygyny (fertilization by a male of multiple females) is likely to be more relevant^77^. In other contexts not explored here (which will also decrease drive strength) pollen limitation can occur^78^. These aspects of the mating system is important to consider since the relative benefit in transmission frequency that *ClvR*-bearing gametes (or those of any other chromosomal TA element) gain in fertilizing ovules, due to loss of competing non-*ClvR* gametes from the same individual, decreases as the number of competing non-*ClvR* gametes from other males increases. Here we provide some representative examples of outcomes when a pan-gamete killing *ClvR* is introduced into a population and the mating system is monogamous—a best case scenario in which *ClvR* gametes from a single male monopolize the ovary of a female—or polyandrous, with five or twenty males each contributing 1/5th or 1/20th of their pollen to a female. We also consider the role of gamete fitness costs, as might arise due to incomplete rescue or cleavage-induced aneuploidy^79^ that manifests itself as death during the haploid stage. The effects of maternal carryover of Rescue activity are also explored.

We first consider population modification. Fig. 5a-c shows examples in which *ClvR* is introduced at a frequency of 10% into a WT population. The LOF allele creation rate is set to 95%, somewhat lower than the rate inferred from the results of our experiments with *ClvR*^ubq7^(Fig. 3 and Fig. 4). Gametic fitness costs (dominant because they are in the haploid stage) were varied between 0% and 15%. Maternal carryover was set to zero as exploration of other scenarios shows it has very little impact on population modification. With monogamous mating *ClvR* spreads rapidly over a range of fitness costs (Fig. 5a). In the presence of polyandry drive is slowed and fails for some higher fitness costs (Fig. 5b,c). However, spread to high frequency can be restored if the introduction frequency is increased (Extended Data Fig. 7). When drive does occur, *ClvR* spreads to allele fixation. This is because whenever a non-*ClvR* homologous chromosome is present with *ClvR* (and the LOF allele creation rate is high) the former has a very high probability of being eliminated from the viable gamete pool since it lacks a Rescue transgene.

**Fig. 5:**
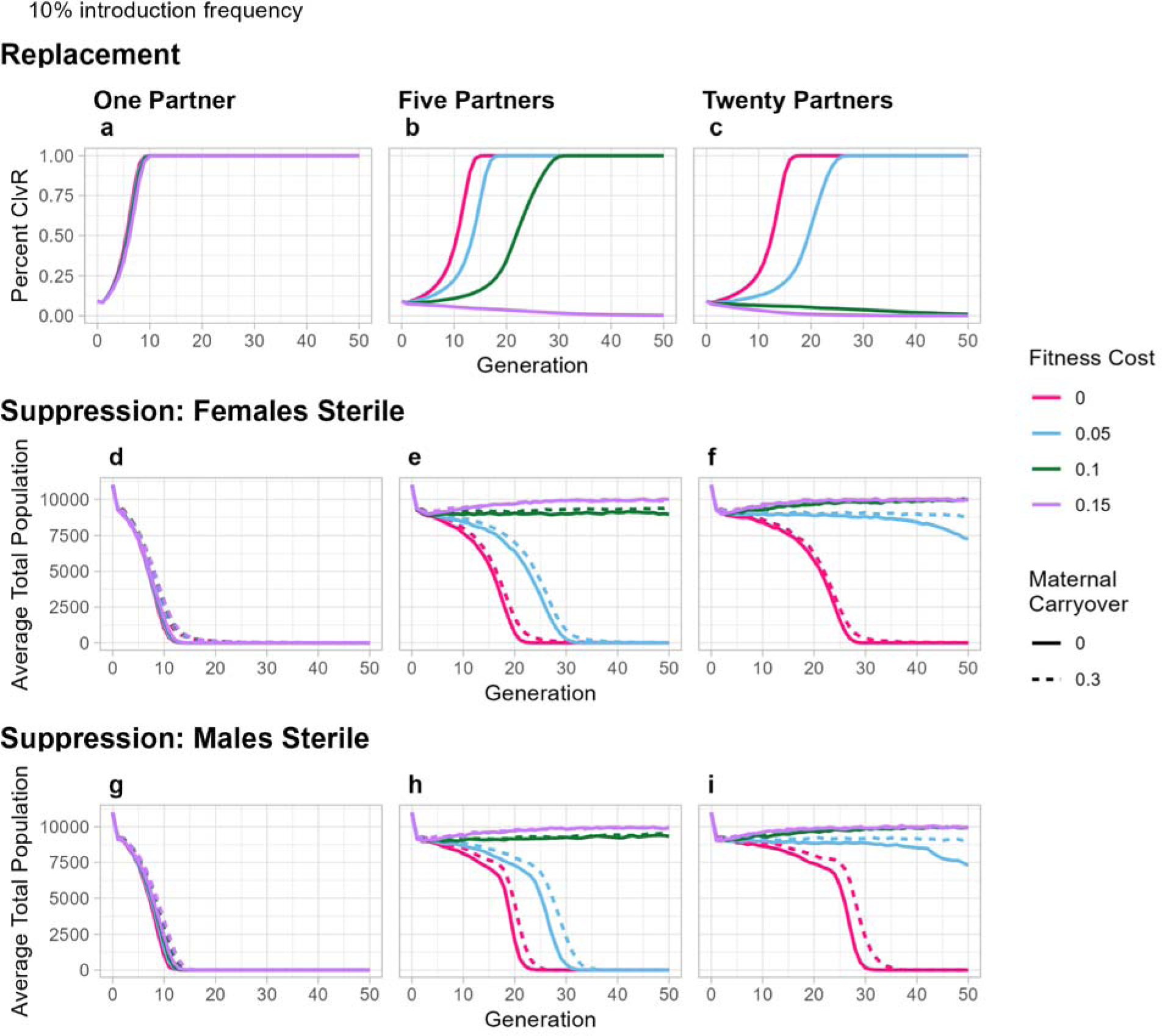
Predicted behavior of *ClvR* for population modification and suppression. **(a-c)** Population modification. *ClvR* is introduced as homozygous males at a frequency of 10% of the starting population, which is at carrying capacity, 10,000 individuals. The mating system is monogamous **(a)**, or polyandrous, with 5 males each providing 1/5th of the pollen needed to fertilize all ovules of an individual female **(b)**, or 20 males each providing 1/20th of the pollen needed **(c)**. Fitness costs are incurred by gametes (a probability of not being able to participate in fertilization, if chosen by the model). Maternal carryover is set to zero. Lines represent the average of 10 runs. **(d-f)** Population suppression with a transgene inserted into a recessive locus required for female sporophyte fertility. *ClvR* is introduced as above, at a frequency of 10%. The mating system is monogamous **(d)**, or polyandrous, with 5 males each providing 1/5th of the pollen needed to fertilize all ovules of an individual female **(e)**, or 20 males each providing 1/20th of the pollen needed **(f)**. Fitness costs are as above. Maternal carryover is set to zero or 30% (the approximate value observed in our experiments with *ClvR*^ubq7^). **(g-i)**. As with **d-f**, but with the *ClvR* inserted into a locus required for male sporophyte fertility. For these simulations homozygous females were released into the population since homozygous males are sterile. For all panels lines represent the average of 10 runs.

For population suppression we consider two scenarios, in which *ClvR* is located in a gene (thereby disrupting it) whose recessive LOF in the sporophyte results in female (Fig. 5d-f) or male (Fig. 5g-i) infertility (as originally outlined in^21^). As above, gamete killing and rescue occurs in both sexes and some level of maternal carryover rescue of LOF allele-bearing gametes may be present. In both scenarios a gamete killer can drive the population towards a homozygous male or female sterile state, resulting in population extinction. A 30% maternal carryover rescue of LOF alleles has a modest negative effect on drive towards a homozygous female sterile state (Fig. 5d-f), while 100% maternal carryover rescue prevents suppression (Extended Data Fig. 8). These negative effects arise because rescue of LOF alleles allows WT alleles at the female fertility locus to persist for some time in non-*ClvR* progeny of a *ClvR*-bearing mother. As with population modification, polyandry and fitness costs can slow or prevent drive and suppression (Fig. 5d-i) but increasing the introduction frequency can be used to compensate (Extended Data Fig. 7).

### *ClvR* drive behavior in response to mutations in cis and trans

TA-based drive depends on the creation of the toxin—in the case of *ClvR* the LOF alleles created by Cas9/gRNAs—and thus is dependent on Cas9/gRNA activity, and the presence of cleavable target sites. As detailed in our earlier work, population modification by *ClvR*s that spread by killing specific zygote genotypes is relatively insensitive to the presence of a high frequency of Rescue/Cargo/gRNA-only alleles lacking Cas9 function^19^. This fact can be utilized to create versions of *ClvR* that show strong, but ultimately self-limited drive for population modification when Cas9 and Rescue/Cargo/gRNA constructs are located at different positions in the genome^38^*. ClvR*-based gamete killers behave similarly. This can be inferred from the results shown in Fig. 6a,b in which a split *ClvR* (Cas9 at one location and Rescue/Cargo/gRNAs at another nearby) is introduced at a frequency of 10%, and Cas9 (which is assumed to be the source of any fitness cost to carriers) recombines away from the Rescue/Cargo/gRNA at a frequency of 1%. Cas9 and gRNAs drive the accumulation of LOF alleles, which select for Rescue/Cargo/gRNAs (Fig. 6a), and initially the tightly linked Cas9, which also increases in frequency (Fig. 6b). However, over multiple generations recombination onto a non-*ClvR* chromosome leads to loss of Cas9 (analogous to a very high mutation rate) when it finds itself alone in LOF gametes lacking Rescue activity. This, in conjunction with Cas9 loss due to natural selection when its presence results in a fitness cost, brings about an eventual end to drive potential (no new LOF alleles can be created), but often not before the combination of intact elements and Rescue/Cargo/gRNA has spread to fixation. A second example that illustrates the resilience of a *ClvR* gamete killer for population modification to loss of Cas9 activity is shown in Fig. 6c,d, in which *ClvR* is introduced at a starting frequency of 10%, with 20% of these elements lacking Cas9 function. Drive of Rescue/Cargo/gRNA-bearing elements (intact elements and Rescue/Cargo/gRNAs) proceeds to allele fixation with only modest delays as compared with a 10% introduction of intact *ClvR*s (Fig. 6c; compare with Figure 5b,c). The frequency of intact elements, which also reflects the frequency of Cas9, initially goes up. However, once the combination of intact elements and Rescue/Cargo/gRNAs reaches fixation the former begins to be eliminated since its presence is associated with a fitness cost that Rescue/Cargo/gRNA elements lack (Fig. 6d).

**Figure 6:**
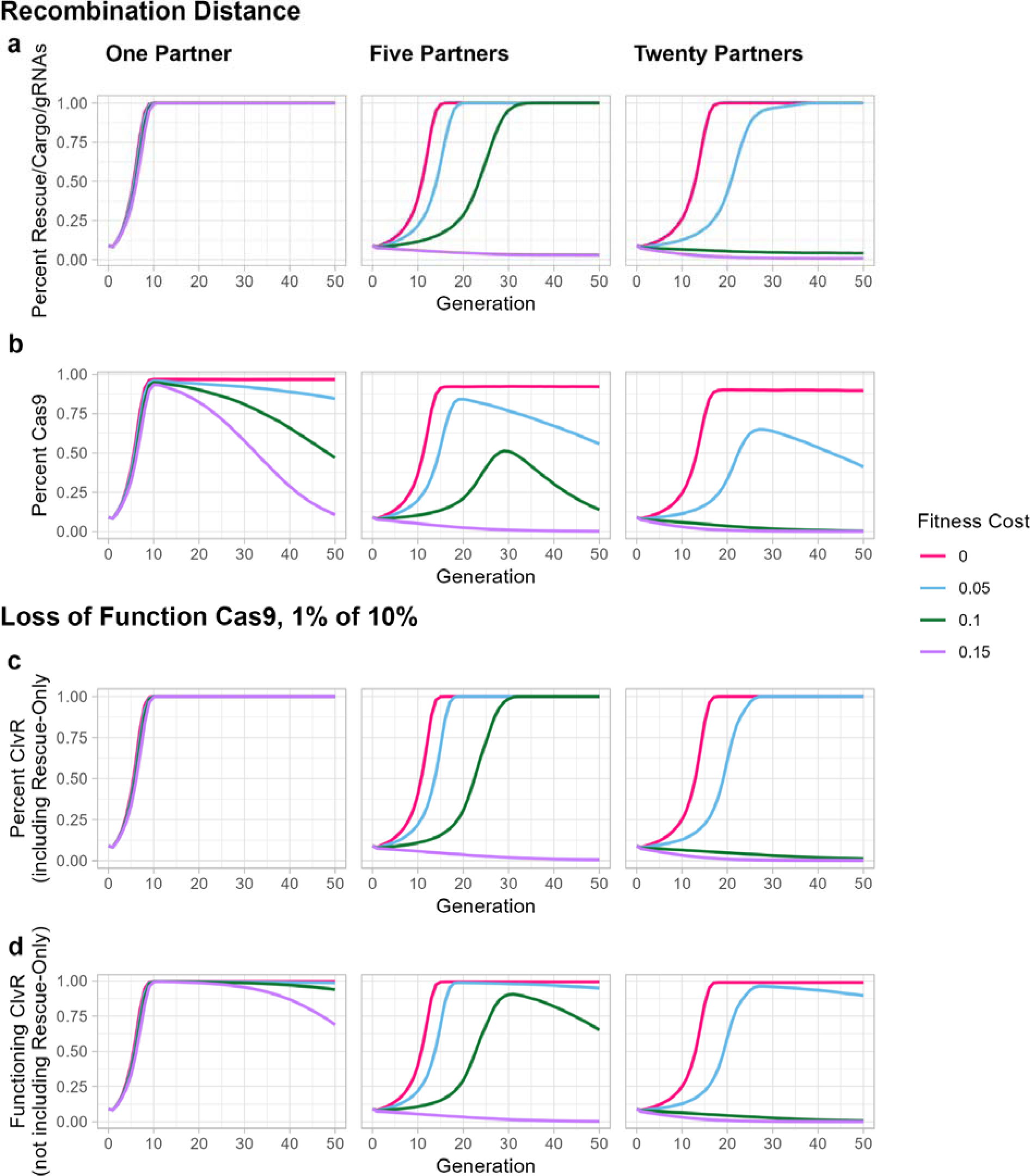
C*l*vR gamete drive for population modification tolerates the presence of significant frequencies of Rescue-only elements lacking Cas9 or gRNA function. **(a-b)** *ClvR* is introduced at an allele frequency of 10%. Recombination distance/frequency between Cas9 and Rescue/Cargo/gRNAs is 1%. **(a)** The frequency of the combined genotypes carrying a Cargo (intact elements and those consisting of Rescue/Cargo/gRNAs) is shown for one, five and 20 mating partners. **(b)** The frequency of Cas9 (which largely reflects its frequency in intact elements) initially increases due to linkage Rescue/Cargo/gRNAs. Recombination into LOF gametes lacking a source of Rescue and natural selection then cause Cas9 to be lost overtime. **(c-d)** *ClvR* is introduced at a frequency of 10%, with 20% (a very high frequency) of these elements lacking Cas9 (Rescue/Cargo/gRNA). **(c)** The frequency of the combined genotypes carrying a Cargo (intact elements and those consisting of Rescue/Cargo/gRNAs) is shown for one, five and 20 mating partners. **(d)** The frequency of intact elements (which also reflects the frequency of Cas9), goes up initially but then falls as Cas9 is lost, leaving the population (when drive is successful) consisting primarily of Rescue/Cargo/gRNA-only elements. For all panels fitness costs are associated with Cas9, and are thus absent in carriers of Rescue/Cargo/gRNA-only elements. Lines represent averages of 10 individual runs.

Drive to allele fixation for population modification can be maintained in the presence of a high frequency of Rescue/Cargo/gRNA-only elements because most of the elements (containing Cas9 and gRNAs) continue to push non-*ClvR* alleles out of the population through creation of LOF alleles. Since this occurs in gametes, not zygotes, the Rescue/Cargo/gRNA-only element can only provide a respite from killing until the non-*ClvR* allele finds itself in a heterozygote carrying an intact *ClvR* element, whereupon it is fated to die in a LOF gamete. Similar considerations will apply to gamete killers that work through a traditional protein-based TA mechanism. Antidote-only alleles can provide a respite from killing. But so long as outcrossing brings the non-element-bearing chromosome into regular contact with intact elements it is fated to be lost. That said, if the presence of the Toxin (or in the case of *ClvR* the Cas9 and gRNAs that create the LOF toxin) results in a cost to carriers then its mutation to inactivity will ultimately lead to a population composed of Cargo-bearing Rescue/Antidote-only elements (Fig. 6d). These, if they also carry a fitness cost and are not already at allele fixation, will ultimately be lost through natural selection, returning the population to a wildtype state, a process originally described in modeling of *Medea*, a maternal-effect zygote killing TA element^16^.

Population suppression by a *ClvR* (or other TA element) inserted into a recessive sporophyte fertility locus is also able to tolerate some level of Rescue/Antidote-only alleles. This behavior is illustrated in Fig. 7 for a population in which a *ClvR* located in a gene required for sporophyte female fertility is introduced at a starting frequency of 10%, with 1% of these elements lacking Cas9 (Rescue/gRNA only elements). Under conditions in which population elimination occurs when an intact *ClvR* is introduced at a frequency of 10% (Fig. 5d-f), elimination also occurs when some Rescue/gRNA-only elements are present. This occurs, as in the case of population modification, because the high frequency of intact elements continues to create LOF alleles, which push the population towards allele fixation for Rescue-bearing elements (intact and Rescue/gRNA elements), a homozygous infertile state for one of the sexes. In contrast, when the fraction of Rescue/gRNA-only elements is 20% of the 10% introduction frequency, or a split *ClvR* is utilized, with a 1% recombination distance between Cas9 and Rescue/Cargo/gRNA, more runs fail (Extended Data Fig. 9). Failures occur in small populations that contain at least one WT allele at the essential gene and fertility loci, and few or no intact elements able to create LOF mutations that select against the WT fertility locus. Other features of the population such as inbreeding and spatial structure can further decrease the probability of successful suppression. Inbreeding decreases the mean fitness of TA element-bearing individuals, while spatially structured populations can allow for local extinction and repopulation with viable and fertile genotypes through migration^80–83^.

**Figure 7.**
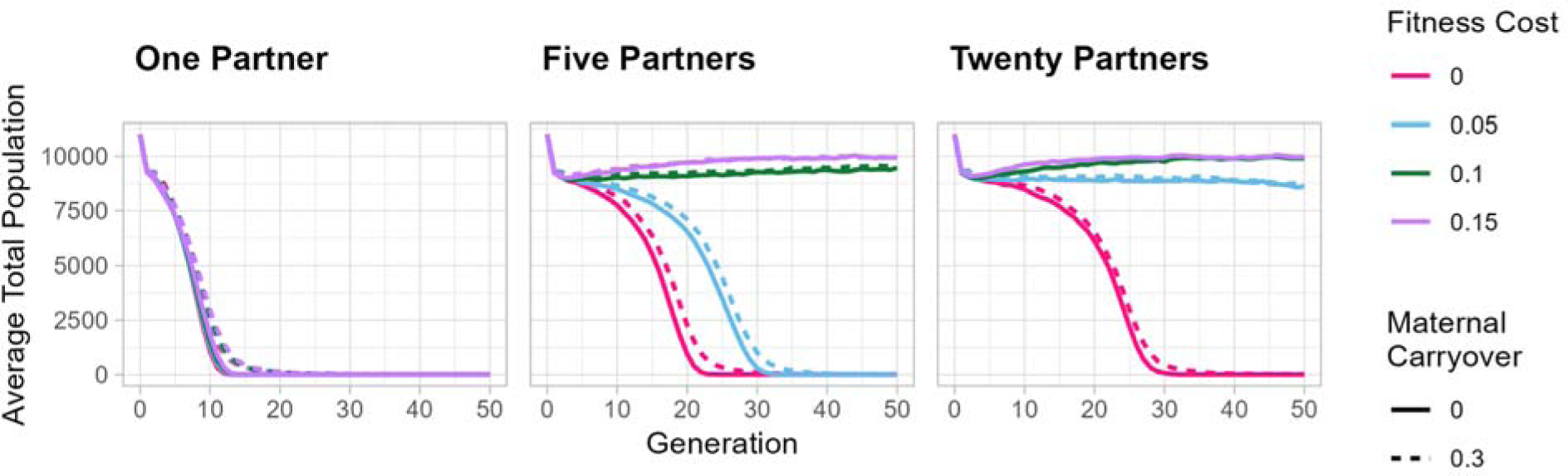
*ClvR* gamete drive for population suppression tolerates the presence of significant frequencies of Rescue-only elements lacking Cas9 or gRNA function. Population suppression occurs when 1% of *ClvR*, located in a female fertility gene and introduced as transgenic males at a frequency of 10%, consists of elements that lack Cas9 (Rescue/gRNA). Outcomes are comparable to those observed when *ClvR* is introduced at a frequency of 10%, and all elements are intact (Fig. 5d-f). Fitness costs are associated with Cas9.

Sequence polymorphisms (naturally occurring or arising through inaccurate DNA repair) that create uncleavable but functional versions (resistant alleles) of a target gene are generally detrimental to gene drive (reviewed in^1,3^). Resistant versions of the essential gene can allow non-*ClvR* chromosomes to survive in gametes produced by *ClvR* carriers. However, selection against the non-*ClvR* allele is still very strong in most gametes, which have or will have LOF alleles in the future. In the context of population modification this can allow *ClvR* alleles to spread to fixation. This is shown in Fig. 8a for a *ClvR* introduction frequency of 10% into a wild population carrying resistant alleles at a frequency of 1% (Compare with Fig. 5a-c). The frequency of the resistant allele increases along with that of *ClvR* (Fig. 8b). In those populations in which the *ClvR* allele ultimately reaches fixation the frequencies of LOF and resistant alleles stabilize because there is no longer a fitness difference between carriers of one versus the other (Fig. 8b). In contrast, when drive does not proceed to allele fixation there is strong selection against the LOF allele, which is then lost along with the *ClvR* allele through natural selection.

**Figure 8.**
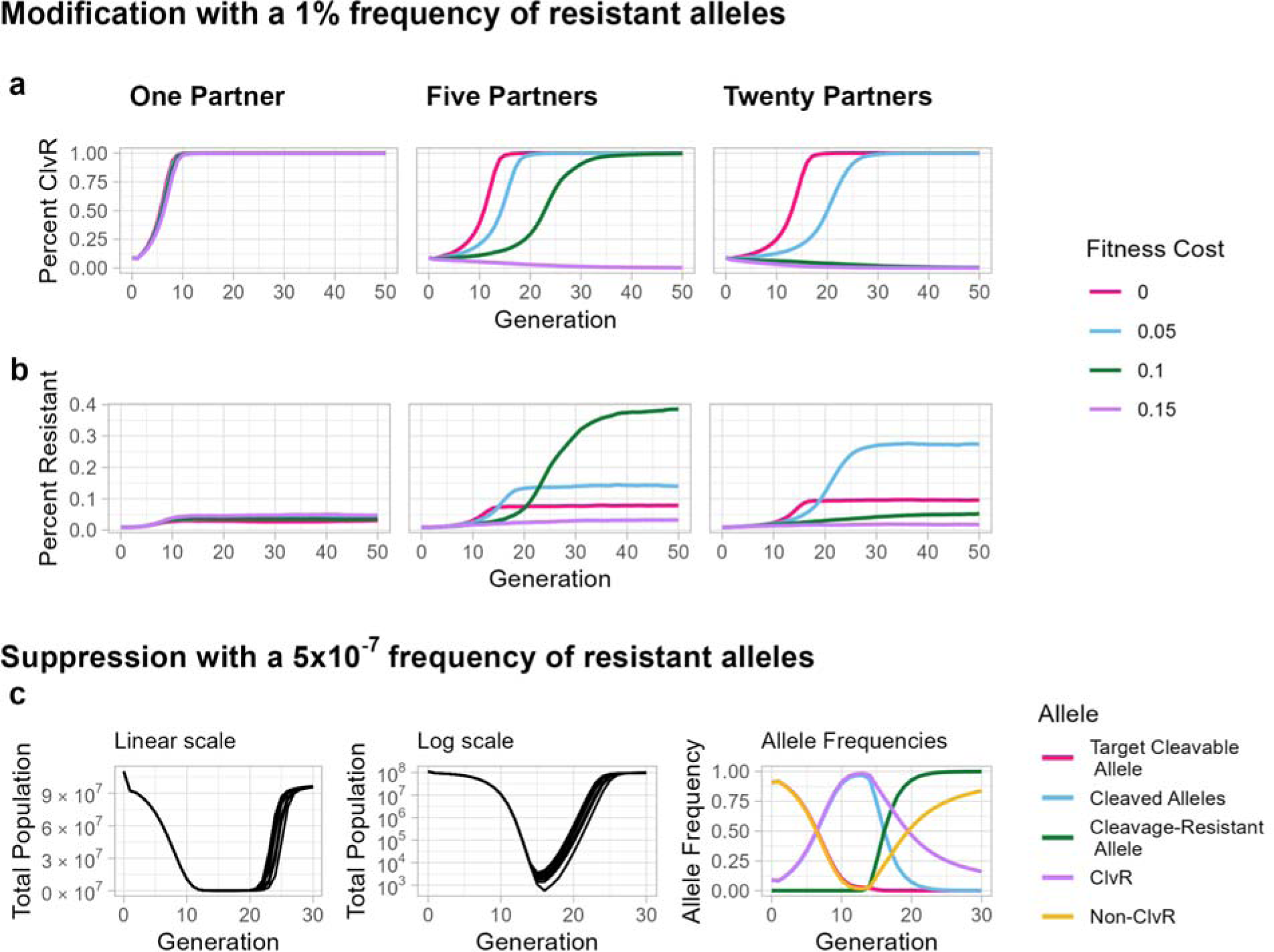
Population modification but not population suppression can occur in the presence of resistant alleles. **(a,b)** Population modification in the presence of a resistance allele frequency in the wild population of 1% and a *ClvR* introduction frequency of 10%. **(a)** *ClvR* spreads to allele fixation under conditions that also support drive when no resistant alleles are present (Fig. 5a-c), but with some delay. **(b)** The LOF alleles created by *ClvR* select for the presence of resistant alleles, which increase in frequency. However, *ClvR* can still reach allele fixation in many cases (panel **a**) because the large number of LOF alleles created from cleavage sensitive essential gene alleles still select against the non-*ClvR* allele. If *ClvR* reaches fixation the frequency of resistant alleles stabilizes because these alleles no longer have a selective advantage. **(c)** *ClvR* is introduced at a frequency of 10%, with a resistance allele frequency in the wild population of 5×10^−7^ (100 resistance allele heterozygotes in a population of 100,000,000 diploid individuals). Total population over time is plotted on a linear scale (left panel) and a log scale (middle panel). The right panel shows the dynamics of alleles at the *ClvR* and essential gene loci over time Finally, when *ClvR* or other TA gametic drive elements are located in a recessive sporophyte fertility gene there are several other possible mechanisms by which population suppression can be defeated. These elements drive against the WT allele at the fertility locus, suppressing the population as sterile homozygotes accumulate. Movement of the TA cassette or the WT allele of the fertility locus to a new location can disrupt this relationship. Thus, if the TA cassette moves elsewhere—through transposition or some other very rare event occurring probably in a single individual— this creates a new allele that can rescue the survival of gametes lacking a TA element at the original location. Because the new TA allele sits at a neutral location it can perform this rescue function without risk of infertility (as would occur in homozygotes for the original TA cassette), thereby preventing population extinction, though not drive. This is shown in Extended Data Fig. 10, in which *ClvR*s are introduced at a frequency of 10%, and 1×10^−5^ of those *ClvR* individuals have one copy of *ClvR* translocated to an unlinked neutral position. An initial population suppression is followed by a rebound. A similar effect is observed when rare individuals in a wild population carry a WT version of the fertility locus at an unlinked position (Extended Data Fig. 10). Since all eukaryotic chromosomal TA elements (as well as other chromosomal drive elements, such as engineered underdominance) spread by driving against counterpart alleles on the homologous chromosome, and drive meant to bring about population suppression strongly selects for suppressor mutations, it will be important to understand the frequency with which genes and multi-gene cassettes move to new locations and/or are present at multiple locations in existing populations.

Resistant alleles pose a much greater challenge to population suppression. *ClvR*-based selection against the non-*ClvR* allele is generally outweighed (in small populations experiencing density-dependent growth) by the large fitness benefit associated with being a fertile heterozygous *ClvR* sporophyte, made possible by the presence of resistant alleles. An example of suppression failure for a *ClvR* introduction frequency of 10%, with 5×10^−7^ of these (100 heterozygous individuals in a population of 100 million) being resistant, is shown in Fig. 8c, left two panels. The population undergoes a large initial drop in numbers. This is associated with an initial rise in the frequency of LOF and *ClvR* alleles (Fig. 8c, right panel). The population then rebounds as resistant alleles and non-*ClvR* alleles, which allow survival and fertility respectively, are selected for and accumulate (Fig. 8c, right panel). Thus, as with other cleavage-based population suppression drives such as homing, prevention of resistant allele formation is essential. Results from work in *Drosophila* on *ClvR* by ourselves and others^19,20,37,39,40^, as well as this study and that of Liu et al in Arabidopsis^59^, along with the results of modeling^84^, suggest that multiplexing of gRNAs (4 or more) may be able to achieve this goal (though see discussion of hybridization below for a context in which this may not be the case).

The above suppression failures due to cassette/gene movement to a new location can be prevented if the TA drive cassette sits at a neutral location and uses a site-specific nuclease to create LOF alleles in the fertility locus (wherever it is located) only after the time during which the gene product is needed for sporophyte fertility, perhaps during meiosis. In this configuration the TA element mediates gametic drive towards homozygosity, while cleavage and LOF allele creation in the fertility gene during meiosis (which must still avoid resistant allele formation) ensures that all homozygous progeny (males or females), but not heterozygotes, are sterile. However, transcriptional regulatory sequences/Cas9 variants that can achieve this goal with high efficiency have yet to be identified.

Finally, we note that in the context of population modification the Cargo will also mutate to inactivity at some frequency in any gene drive system. It may also lose effectiveness due to evolution of the host or pathogen it is meant to counter. Recombination between chromosomal TA drive elements and WT homologous chromosomes can also—in some configurations, but not others^19,34^—lead to the creation of Rescue-only elements lacking Cargo. For chromosomal TA drive elements the consequences of Cargo loss/failure can be ameliorated by carrying out a second round of population modification in which a first-generation element, at its original location, competes against a next-generation element (carrying a new Cargo, Toxin and Antidote, and a copy of the first-generation Antidote) at the same location. The latter drives itself in while driving the first generation element (and any remaining WT alleles) out of the population^34,37^. Next-generation gamete killers are expected to be particularly efficient since they spread to allele fixation, which leads to complete elimination of first-generation elements.

## Discussion

Our results, along with those of Liu and colleagues^59^ argue that gamete killers based on a *Cleave* and *Rescue* mechanism provide a general strategy for achieving gene drive-mediated population modification or suppression in outcrossing diploid plants. *ClvR* elements utilize a simple toolkit of components that should be available in many species: a site-specific DNA-modifying enzyme such as Cas9 and the gRNAs that guide it to specific targets, sequences sufficient to direct gene expression in cells that will become the germline (which, as in our work, need not be germline-specific), an essential gene to act as target, and a recoded version of the essential gene resistant to sequence modification and able to rescue the LOF condition. For population modification these components are best located at sites distant from the target essential gene. For several strategies for population suppression the element needs to be located in a gene whose recessive LOF in the sporophyte results in either male or female infertility. Many such genes are known^85–88^, particularly for male fertility. Alternatively, if LOF alleles of the sporophyte fertility gene can be efficiently created late in germline development, after the time when gene function is needed for fertility, then the element can be located anywhere in the genome.

An alternative strategy for population suppression in some plants is suggested by modeling and experiments in animals focused on the creation of sex-linked gamete killers. The goal of these efforts is to create Y-linked killers of X-chromosome-bearing sperm, resulting in males that only produce male progeny (reviewed in^1,3^). Such a system can be used to drive population suppression or elimination (towards an all-male state) when sperm is not limiting and females mate with one or a few males^13,42^. Many dioecious plants lack sex chromosome with well-defined regions whose presence is sufficient to confer a specific sex on carriers. However, for those that do carry such regions^89,90^ it may be possible to engineer a similar behavior. For example, if a pan-gamete killing *ClvR* (or a killer of non-*ClvR* gametes only in pollen) is tightly linked to a gene or genes that are sufficient for male sex determination (e.g.^91,92^) then *ClvR*-bearing individuals (by definition males) will only produce *ClvR*-bearing sons and pollen that gives rise to male progeny. As discussed in the context of Fig. 5, drive towards population extinction becomes weaker as the level of polyandry increases, but may still occur in a timely manner if the introduction frequency is increased. Notwithstanding these points it is important to note that even in outcrossing species inbreeding and spatial structure can work to prevent elimination of a population ^80–83^.

Finally, the *Cleave* and *Rescue* mechanism could also be used in a non-gene drive strategy for population suppression. Modeling and experiments in insect systems show that periodic releases of males carrying an autosomal transgene that gives rise to fertile males and inviable or sterile females can bring about population suppression or elimination by driving a progressive decrease in the number of fertile females^93–95^. Such an element does not show self-sustaining drive because it finds itself in dead-end females half the time. However, its persistence over multiple generations in fertile males provides an ongoing force that contributes to a reduction in the number of fertile females. Such an element could be created in diploid plants (dioecious, monoecious or hermaphrodite), though self-fertilization will always reduce effectiveness in hermaphrodites and monoecious species that lack strong incompatibility systems. There are several possible approaches. In one a pan gamete *ClvR* (or other protein-based TA element) is located at a neutral position and carries a transgene that dominantly blocks female gamete development. However, this system is not evolutionarily stable since if the transgene needed to block female fertility is inactivated by mutation one is left with a self-sustaining gamete killer drive element. A more stable strategy involves locating a pan gamete killing *ClvR* element within (thereby disrupting) a gene whose expression in female gametes is required for their survival. Reproductive structures of individuals carrying this construct only produce *ClvR-*bearing pollen. Female gametes that inherit the *ClvR* die because they lack the female gamete essential gene while those that lack the *ClvR* die because they lack a functional copy of the pan gamete essential gene targeted by *ClvR* for LOF allele creation. Loss of Cas9 function through mutation in a small fraction of the suppression strain can allow the survival of some Rescue/gRNA-only individuals, but their presence does not block suppression because they cannot be transmitted through the female germline.

Any gene drive method, when it does not provide an unalloyed fitness benefit to carriers, is sensitive to mutational inactivation. Second site suppressors may also be selected for that block drive or its intended consequences. Our modeling suggests that population modification is relatively insensitive to mutation of Cas9/gRNA to LOF or the presence of a modest frequency of resistant alleles at the essential gene locus. Population suppression can also occur in the presence of a modest frequency of elements that lack Cas9 but is very sensitive to the presence of resistant alleles, as with homing based strategies. Suppression through some mechanisms is also sensitive to movement of the TA element or a WT allele of the fertility gene to a new chromosomal location. Finally, while the Cargo can also undergo mutational inactivation or loss of efficacy, population modification and suppression strategies with chromosomal TA elements can be made resilient—able to recover from breakdown—using next-generation elements that drive old elements out of the population while driving themselves in.

Our experiments focused on cleavage and rescue of the ubiquitously expressed R-SNARE YKT61 gene. It is likely that many other ubiquitously expressed housekeeping genes can be targeted to similar effect. Alternatively, drive can be limited to one sex or the other by targeting genes required more specifically for gametogenesis in only one sex (e.g.^49,50^). Liu and colleagues used just such an approach, targeting the No Pollen Germination 1 gene for cleavage and rescue in *Arabidopsis thaliana*. Carriers of this construct show high levels of *ClvR*-biased segregation distortion through pollen but not ovules ^59^. Ideally Cas9 expression, cleavage and LOF allele creation would be limited to cells of the appropriate reproductive organ or meiosis, so as to minimize fitness costs associated with Cas9 expression or heterozygosity for LOF mutations in the target gene (haploinsufficiency), and allow for targeting of fertility genes after the time in development they are needed. These were the reasons we tested regulatory sequences from genes with restricted expression patterns that include the future germline: DMC1 and APETELA1, CLAVATA3 and AGAMOUS. Among these only APETELA1 sequences showed evidence of strong drive in males. Low levels of drive were observed in females. Based on the results of experiments discussed above we speculate this may be because of particularly strong maternal carryover rescue of gametic LOF alleles. Even when using the UBIQUITIN10 sequences to drive Cas9 expression, providing many opportunities throughout development to cut and create LOF alleles, we observed a low frequency (∼1%) of uncut/unmodified alleles at the YKT61 locus in escapers. This is not due to transgene silencing since the construct was not present in escaper seeds (Extended Data Fig. 6). Nucleosome structure has been shown to inhibit Cas9 cleavage efficiency^101–103^, and could play a role, though it is surprising that all four sites remained uncleaved. Only the results of more experiments with diverse promoters, and other RNA-targeted DNA sequence modifying enzymes that cleave or create LOF mutations through other mechanisms such as base editing, acting on YKT61 and other target genes, will provide guidance on how best to ensure that all target sites are modified. Regardless, our modeling shows that a low frequency of WT escapers (which are still subject to cleavage in future generations) does not prevent population modification or suppression.

Plants are the backbone of life on earth and the source, directly or indirectly, of most human food. Given this it is important to consider if gene drive in plants constitutes a dual use research of concern (DURC). In the context of biology DURCs are research products that, while designed to provide a clear benefit, could potentially be misused to threaten public health and safety, agricultural crops and other plants, animals, or the environment^96^. As discussed in earlier work^97^, gene drive in plants and animals in general does not lend itself to DURC applications. First, spread of a drive element to high frequency is very slow, because it requires many generations of outcrossing. In the case of plants generations tend to be seasonal (yearly) at best, and inbreeding, which slows drive, is common. Drive elements that work indirectly, by killing those who fail to inherit them, are (as compared with drives that home at high frequency) particularly slow to spread when introduced at modest frequency^1,17,19,20^. Second, in modern agriculture crop breeding, seed production and distribution typically occur under tightly controlled conditions using specific genotypes, making it unlikely other genotypes could be introduced into the production/food chain without detection^98^. Related to this last point, gene drive is easily detected if searched for, either through observation of phenotypic changes in a population or through genome sequencing. Finally, gene drive for population suppression can and has been blocked—many times—through the creation of resistant alleles (reviewed in^1,3^). It has also been blocked through introduction of a transgene that inhibits Cas9 function^99,100^, and it can in principle be blocked using a second modification drive that actively targets key components of the initial drive element of concern. The consequences of population modification (though not necessarily a rapid return to the pre-transgenic state) can also be prevented through the use of next-generation elements that drive a first generation element out while driving itself in^37^.

The above points argue that gene drive in plants is unlikely to constitute a DURC technology. However, the frequent ability of plants to hybridize across species barriers^104,105^ calls attention to several competing challenges related to drive, resistance to drive and gene flow. Gene drive with the *ClvR* system can be limited to a specific species by designing gRNAs that are species specific. If hybridization does occur in this context the WT essential gene alleles from the relative are. by definition, resistant. These will block spread in the non-target species but may still allow population modification in the target species, depending on the rate of hybridization, and thus the frequency of resistant alleles (Fig. 8a,b). However, as noted above (Fig. 8c-e) resistant alleles would prevent population suppression in the target species. Alternatively, gRNAs can be utilized that target the essential gene for LOF allele creation in all possible hybridization partners. This should support population modification and suppression in the target species but may also result in modification or suppression in non-target species. Protein-based TA systems, which typically target conserved biological processes rather than specific genomic sequences, may behave similarly. Thus, in considering TA-based gene drive in plants it will be important to understand the full spectrum of mating partners and possible ecological outcomes associated with drive, both within a target area and in non-target areas connected by migration.

Finally, while our experiments and modeling focused on *Arabidopsis thaliana*, a diploid with a relatively small genome, many plants of interest are polyploid^106^. Large genomes and polyploidy create several challenges. First, large genome size means Cas9 must sample a much larger genomic sequence space in a timely manner^107^, which will require increased expression levels or the use of variants with increased catalytic activity. Second, polyploidy may release duplicated genes, even those encoding highly conserved housekeeping genes, from selective pressures that constrain their coding sequence, making it more difficult to identify gRNA target sites that remain unchanged. The design of gRNAs will be particularly challenging in allopolyploids, which have two or more complete sets of chromosomes from different species. The gene dosage needed for rescue (one or multiple copies) also needs to be explored for polyploids. Suppression mechanisms that require insertion of *ClvR* into a gene required for gamete function may be challenging for related reasons. In sum, while our work and that of Liu et al^59^ show that the conditions for *ClvR*-based gene drive in plants can be achieved, much remains to be understood as to species and contexts in which the key mechanisms required for drive (high frequency creation of LOF alleles and rescue) are most likely to be efficient and evolutionarily robust, and in which gene flow can be managed.

How might gene drive be applied in plants? Spread of agricultural traits, weed control and evolutionary rescue have all been suggested^4–6^. As discussed above in the context of DURC research, drive itself is unlikely to be used in spreading agronomic traits into major production crops since seed production (often of hybrids) and distribution is a highly regulated process^98^. Gene drive that causes death of non-carriers (as compared with homing, which can immediately create homozygotes from heterozygotes) is also unlikely to dramatically speed the breeding process. Possible exceptions where population modification could provide some utility include wild plants used as forage for livestock on the range, or in aquafarming. In these contexts, potential target species will often not have undergone selection by humans, and thus might benefit from introgression of genetic changes that enhance food traits and/or resilience in the face of current or impending environmental stresses.

The most proposed application of gene drive in plants is weed control. This could take the form of population suppression or sensitization, in which the goal is to drive a trait into the population that makes the species less fit in a managed agricultural environment or specifically sensitive to some other intervention, such as herbicide application. One species that has been suggested as a good target for gene drive is *Amaranthus palmeri* (Palmer amaranth)^4^, an invasive agricultural weed that is very economically destructive and difficult to manage^108^. Features that make Palmer amaranth amenable to gene drive-mediated suppression and/or sensitization are that it is an annual, dioecious, a diploid, and a region containing genes associated with male sex determination has been identified^91,109^. Finally, in many locations Palmer amaranth has become resistant to available herbicides, with a key source of resistance (and thus a good target for mutation) being a large autonomously replicating extrachromosomal circular DNA transmitted through pollen^110,111^. These attractive features for drive that results in suppression and/or sensitization notwithstanding, Palmer amaranth also exemplifies potential challenges and tradeoffs. It produces a very large number of seeds (100,000-500,000) per plant and can also hybridize with related species^112^, some of which are also weeds. It is also native to Northern Mexico and the Southwestern United states and has cultural significance to Native Americans, who have used it as a food source^108^. These facts highlight the topics of evolutionary stability, gene flow and social acceptability, subjects that have not been explored in plants, particularly in the context of highly managed modern agricultural environments where the goal will be very local rather than global population control.

Finally, gene drive in plants, as well as animals, has been suggested as a tool for bringing about evolutionary/genetic rescue, the process by which a species threatened with extinction adapts rapidly enough to survive. Evolutionary rescue involves bringing new individuals into a population. This increases population size, buffering it against stochastic fluctuations, while at the same time introducing genetic variants that can decrease inbreeding depression and—when they are present at high frequency— increase absolute population fitness (the ability of the population to increase in size) through adaptation. Here we focus on the role of new adaptive variants. The question evolutionary rescue strategies face is whether modest introductions of these variants can bring about an increase in population fitness before stochastic effects take the population below a critical density that leads to extinction^113,114^. While introduced beneficial alleles will spread through natural selection, the rate of spread (and thus the time the population spends near the critical density) depends on the strength of selection and whether the beneficial alleles are dominant, additive, or recessive. We speculate that there may be some contexts in which gene drive can increase the rate of allele spread, thereby keeping average absolute population fitness (and thus population size) higher than it would be otherwise, supporting recovery. However, modeling that tests this hypothesis by comparing the rescue effects of a beneficial Mendelian allele that spreads through natural selection—which requires the death of non-carriers—with that of a similar allele also subject to a gene drive that does not require the death of non-carrier adults remains to be carried out. Finally, we note that gene drive, but not Mendelian transmission and natural selection, could be also used in an anticipatory manner, to spread genetic variants that do not confer a strong benefit now, but that will be beneficial under likely future conditions.

## Methods

### Synthesis of *Arabidopsis ClvR* constructs

In this study, constructs were assembled using Gibson cloning^115^. The gRNA cassette, composed of four repeats of gRNA with U6 promoters, was cloned with Golden Gate assembly. Enzymes utilized were obtained from NEB, and cloning as well as DNA extraction kits were sourced from Zymo. The *A. lyrata* Rescue gene was synthesized by Twist Bioscience.

We began with an intronized Cas9 variant known as zCas9i (pAGM55285, which was a gift from Sylvestre Marillonnet, Addgene #153212)^116^. In this Cas9 version, we replaced the RPS5 promoter with that of DMC1. Additionally, immediately upstream of the start codon, we incorporated 21 base pairs from AT1G58420, which had previously been demonstrated to enhance translational efficiency^117^.

Finally, we integrated the recoded *A. lyrata* Rescue into the construct. A detailed sequence map is provided in Supplementary file 1.

### gRNA design and cloning

To assemble the gRNA cassette we used the shuttle vectors from Stuttmann et al (pDGE332, pDGE333, pDGE335, and pDGE336, which were a gift from Johannes Stuttmann, Addgene #153241, 153242, 153243, and 153244)^118^. Guides were designed in Benchling to target exon 1 (gRNA1 and 2), exon 2 (gRNA3) and exon 5 (gRNA4) of YKT61 and cloned into the Bbs1 digested shuttle vectors with annealed primers. The final Golden Gate assembly was performed with the Cas9-Rescue plasmid from above and Bsa1.

### Cas9 promoters

We could not detect any cleavage with the intronized Cas9 and the DMC1 promoter, as inferred by the Mendelian inheritance of the full *ClvR* construct in multiple transgenic lines. Based on these results we replaced the intronized Cas9 with one that had no introns but retained the NLS sequences at the termini (Cas9 without introns from pTX168, which was a gift from Daniel Voytas, Addgene #89257)^119^. Additionally, we introduced a mutation (K918N) in the Cas9 sequence that was shown to enhance its catalytic activity^120^. However, the DMC1-Cas9 version without introns also showed no evidence of cleavage. Based on the results obtained with other promoters (APETELA and UBIQUITIN 10) and this version of Cas9, we inferred that the DMC1 promoter is likely to be relatively weak. For all additional promoters tested here we used the version of Cas9 version lacking introns and carrying the K918N mutation.

Using Gibson assembly, we replaced the DMC1 promoter with transcriptional regulatory sequences from APETALA1, CLAVATA*3*, and AGAMOUS (chosen based on their efficacy in previous work utilizing a Cre/Lox reporter)^121^. Finally, we also built two versions of *ClvR* utilizing regulatory sequences that drive more ubiquitous expression, from the UBIQUITIN10 gene and the CaMV35S promoter. Whole plasmid sequencing was performed by Plasmidsaurus, using Oxford Nanopore Technology with custom analysis and annotation. Genbank files of all *ClvR* constructs with attached Gibson cloning and sequencing primers utilized in this study are in Supplementary File 1.

### Arabidopsis handling

All plants in this study were grown in soil with a 16/8 hour light dark cycle. Temperature was 25L. All seeds were planted directly in soil and stratified at 4C for 3 days. Transgenic plants were maintained in separate dedicated room in a hood mounted with a screen and surrounded by sticky tape, to minimize airflow around the plants and to prevent potential insect-mediated pollen movement. A floor-mounted sticky surface surrounding the hood performed a similar function. Transgenic plants were disposed of following autoclaving.

### Arabidopsis Transgenesis

We used the floral dip method with agrobacteria as described previously^122^. *ClvR* plasmids were transformed into GV3101 ElectroCompetent Agrobacterium strain from Intact Genomics. T1 seeds were screened for the FAST red seed marker^123^ and planted as described above.

### Crosses to determine *ClvR* drive activity

Red T1 seeds were grown and allowed to self cross. Siliques (seed pods, with each pod representing the fertilized ovules of a single ovary/flower) of these plants were screened for the FAST red marker again. We looked for plants that showed 100% *ClvR* bearing seeds, suggesting drive activity (Mendelian genetics would result in 75% red seeds). At this stage we saw that *ClvR*^dmc1^*, ClvR*^agamous^*, ClvR*^clavata3^ had less than 100% red seeds in the self crosses and decided not to further characterize these lines.

T2 *ClvR* seeds were grown and pollen from these plants was used in an outcross to WT females to generate heterozygous *ClvR/+* T3 seeds. T3 seeds were grown into adults again to set up reciprocal crosses with WT. ♀*ClvR*/+ for each line were crossed to ♂WT and ♂*ClvR*/+ were crossed to ♀WT (4 crosses per plant, 4 plants per line). Siliques of these crosses were scored for the *ClvR* marker (results in Fig. 2).

Next, we took T4 seeds from individual T3 crosses and repeated a set of reciprocal (male *ClvR/+* to female WT and female *ClvR/+* to male WT) crosses. We crossed 4 plants with 4 crosses per plant for each of 3 individual T3 crosses (results in Fig. 3). We also collected leaf tissue from T4 heterozygous *ClvR* plants and escapers from the ♀*ClvR*/+ X ♂WT cross (non-*ClvR* bearing seeds) to extract DNA and sequence the YKT61 target sites (see below). Sequencing results are in Extended Data Table 1. Finally, we grew T5 escaper seeds coming from ♀*ClvR*/+ and from ♂*ClvR*/+. Leaves of young plants were again collected, and target sites sequenced as described below.

### Molecular analysis of cleavage events

DNA from candidate plants was extracted from leaves with the Zymo Quick-DNA Plant/Seed miniprep kit according to the manufacturers protocol. The YKT61 target region was PCR amplified using primers ykt-cleaveF1 (TAGCATCTCCGAGTAAGGAATC) and ykt-cleaveR2 (CTTATAGATTTAGTTTCCTTTTTTCCCTGT). The PCR fragment was purified following agarose gel electrophoresis and sequenced by Plasmidsaurus at ∼1000X coverage. The resulting raw reads were mapped to the YKT61 reference using minimap2^124^. The alignment file was sorted and indexed with samtools. The output file variants were then clustered with a Python script from Pacific Biosciences (https://github.com/PacificBiosciences/pbampliconclustering). Mutations were analyzed in the output “variantFraction” file. Results are summarized in Extended Data Table 1. Sequencing files are in Supplementary File 2

### T-DNA insertion

To determine the T-DNA insertion site of line *ClvR*^ubq7^ we extracted genomic DNA from two different plants and constructed sequencing libraries using NEBNext® Ultra™ II FS DNA Library Prep Kit for Illumina (NEB #E7805) following manufacturer’s instructions. The libraries were sequenced on Illumina NextSeq2000 in paired end mode with read length of 150 nt to the sequencing depth of 35 million paired end reads per sample. Base calls were performed with DRAGEN BCL Converter and structural variant analysis was performed with the DRAGEN Germline pipeline v3.10.12 against the *Arabidopsis* TAIR 10 genome^125^. The resulting VCF files contained information on large structural variants (insertions). We identified a single T-DNA insertion on chromosome 3 (Chr3:10231731). The insertion was confirmed with PCRs followed by sequencing of the resulting amplicons. Extended Data Fig. 11 shows the genomic location.

### Modeling

Modeling was performed using a stochastic agent-based model with discrete generations written in python. The model uses various classes to keep track of Haploid gametes, Diploid individuals, and Simulation parameters. The lowest level class is Diploid, which tracks the genotypes and alleles of a single individual. The next highest class is Haploid, which tracks its own alleles and also its Diploid parent. The highest class, in which most of the data is stored, is StochasticSim, or FastStochasticSim. This class object contains all the parameters necessary to run a simulation, and can store the individuals that are created over the course of a simulation. The parameters stored by StochasticSim include the alleles and genotypes possible, the haploid and diploid fitnesses associated with each genotype, the fecundity of each individual, and other parameters used over the course of the simulation. The function stochastic_sim calls on a StochasticSim object to perform the simulation and populate the generations. Each generation starts with a pool of adult individuals (the introduced individuals for generation 0, and the previous generation’s mature adults for each following generation). Each adult produces a pool of gametes, and each pool of ovules is matched with one or more pools of pollen, which are congregated together to form one single pollen pool. These gamete pools are then reduced by their haploid fitness costs, which are set as part of the simulation parameters. Surviving pollens and ovules are then matched, with each ovule-pollen pair producing a possible offspring. All possible offspring from these matings are randomly grouped as male or female, and are then culled based on diploid fitness and the density-dependence function *S(P) = g / [ 1 + (g-1)*P/K]* where *P* is the parent generation’s population size, and *K* is the carrying capacity. *S* ranges from some growth factor *g* = 6 at low densities, to 1 at densities near carrying capacity. The offspring that survive the culling become the parents of the next generation. A diagram of this process and more details are included on the Github, and in Extended Data Fig. 12.

The function stochastic_sim is called by run_stochastic_sim, which handles doing multiple runs and writing the data out to files. More details are provided on the specifics of stochastic_sim below.. For the data shown we assumed *ClvR* and the target gene were unlinked, that our *ClvR* element had 95% efficiency in creating LOF alleles, and that the population had a low-density growth rate of 6. More information on the model, the scripts and parameters used to generate the data, and the data itself can be found on https://github.com/HayLab/Pigss

## Data Availability

All data is available in the main text or the supplementary information files. Illumina sequencing reads were deposited to SRA (bioproject: PRJNA1074841)

## Acknowledgements

We thank Elliot Meyerowitz and members of the Meyerowitz Lab Paul Tarr and Carla de Agostini Verna for introducing us to techniques for *Arabidopsis* maintenance, transgenesis, and crossing.

## Contributions

Conceptualization, G.O., T.I. and B.A.H.; Methodology, G.O., T.I., M.L.J., I.A., and B.A.H.; Investigation, G.O., M.L.J, I.A., B.A.H.; Writing – Original Draft, G.O. and B.A.H.; Writing – Review & Editing, G.O., T.I., M.L.J. and B.A.H.; Funding Acquisition, B.A.H.

## Ethics declaration

The authors have filed patent applications on *ClvR* and related technologies (U.S. Application No. 15/970,728 and No. 16/673,823).

---

**Extended Data Fig. 1:**
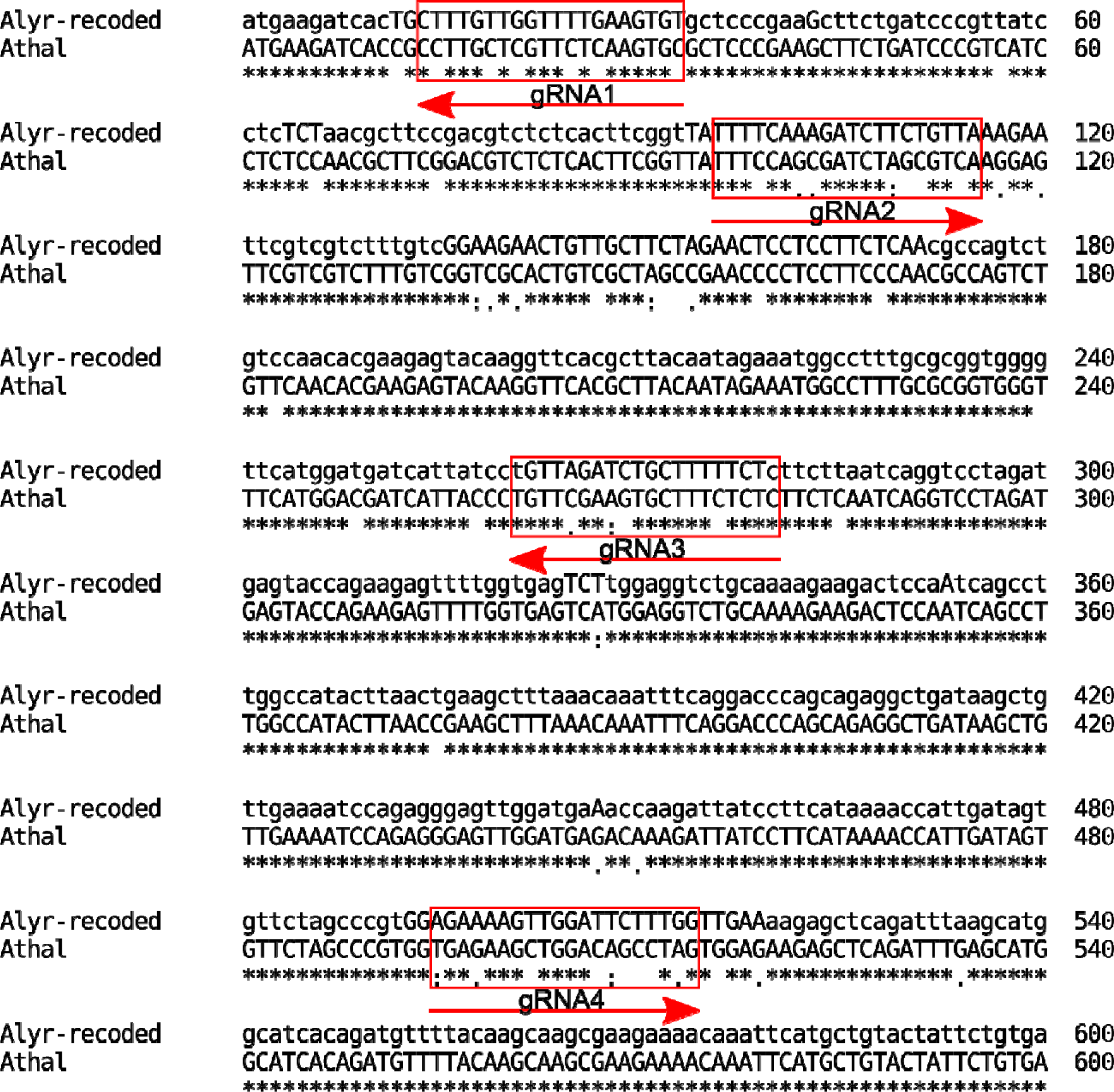
Alignment of the recoded *A. lyrata* rescue coding region to the *A. thaliana* target. Guides are indicated as red arrows. Note that the full sequence of the *A. lyrata* YKT61 genomic region used for rescue (Supplemental File 1) contains many additional differences from the equivalent *A. thaliana* sequence, in regulatory sequences, introns and 5’ and 3’ UTR. The amino acid sequences of the two proteins are identical.

**Extended Data Fig. 2:**
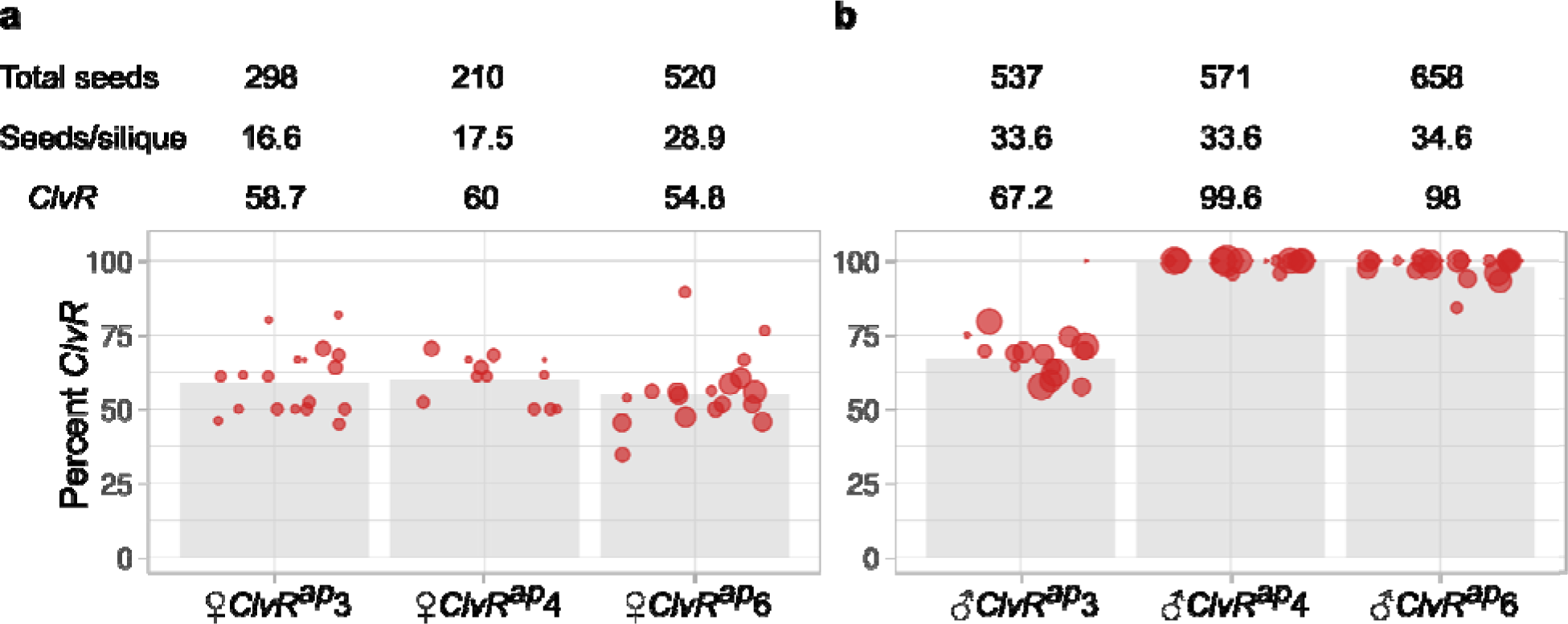
T3 heterozygous *ClvR* crosses for (a) female *ClvR^ap^* and (b) male *ClvR^ap^*. T3 *ClvR*^ap^/+ heterozygotes were grown to adulthood and their ovules (left three columns) or pollen (right three columns) used in outcrosses to WT. Bar graphs show the number of siliques scored (red circles) and the percent *ClvR* seeds produced in the T5 generation. The number of seeds within each silique scales with circle size. Counts are in Supplementary Table S2.

**Extended Data Fig. 3:**
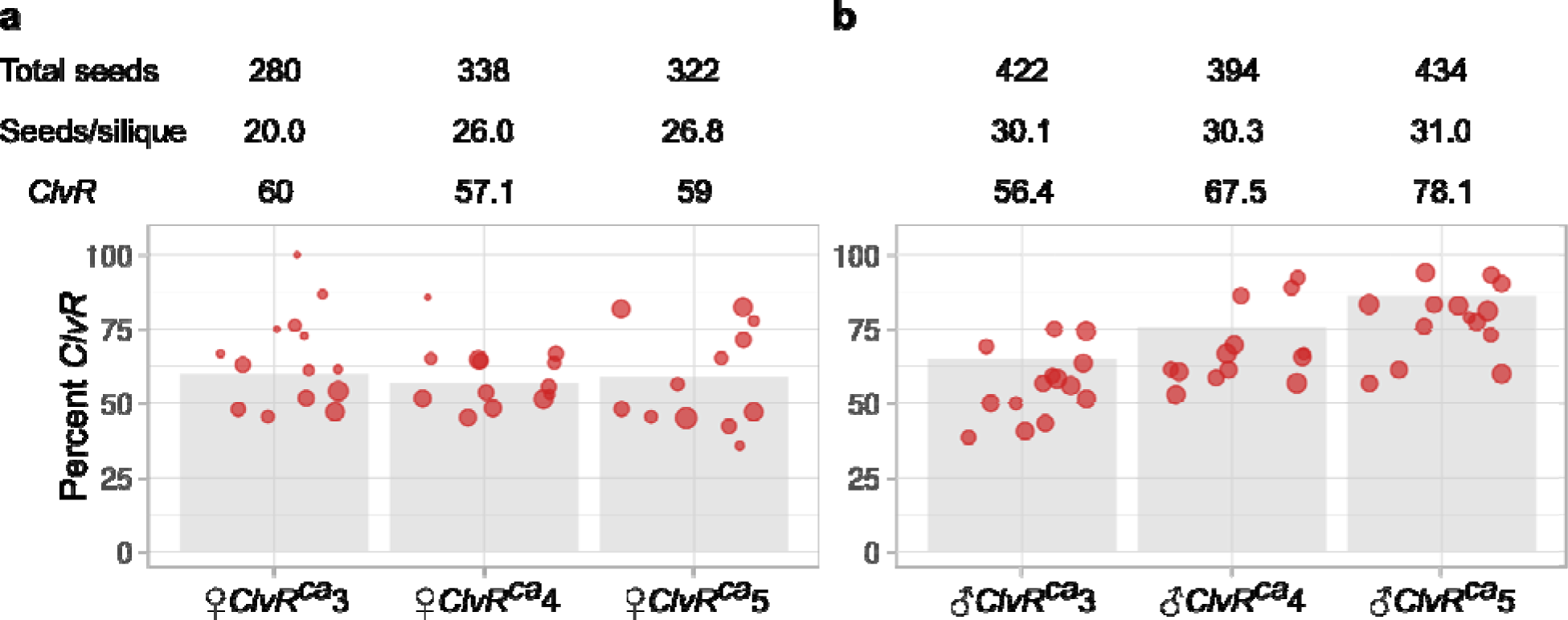
T3 heterozygous *ClvR* crosses for (a) female *ClvR^CaMV35S^* and (b) male *ClvR^CaMV35S^*. T3 *ClvR^CaMV35S^/*+ heterozygotes were grown to adulthood and their ovules (left three columns) or pollen (right three columns) used in outcrosses to WT. Bar graphs show the number of siliques scored (red circles). The number of seeds within each silique scales with circle size. Counts are in Supplementary Table S3.

**Extended Data Fig. 4:**
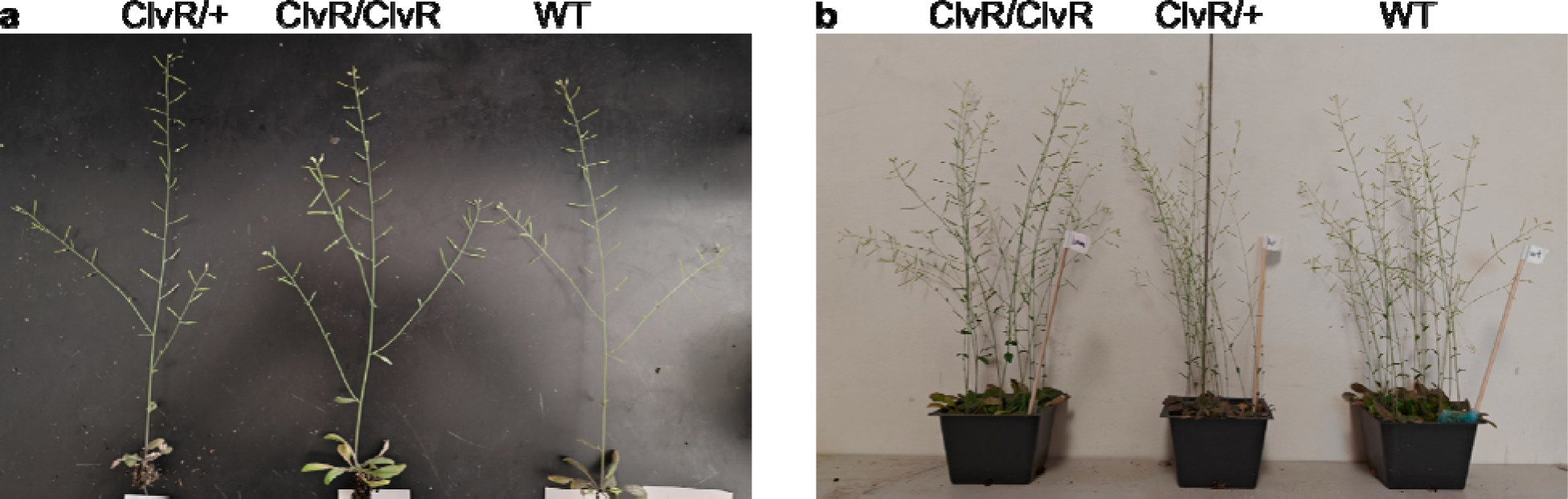
Images of individual (a) and whole pots (b) of heterozygous *ClvR^ubq^*, homozygous *ClvR^ubq^* and WT plants. In **a** individual plants have been removed from their pots and laid flat against a black background. **b** shows pots containing multiple plants.

**Extended Data Fig. 5:**
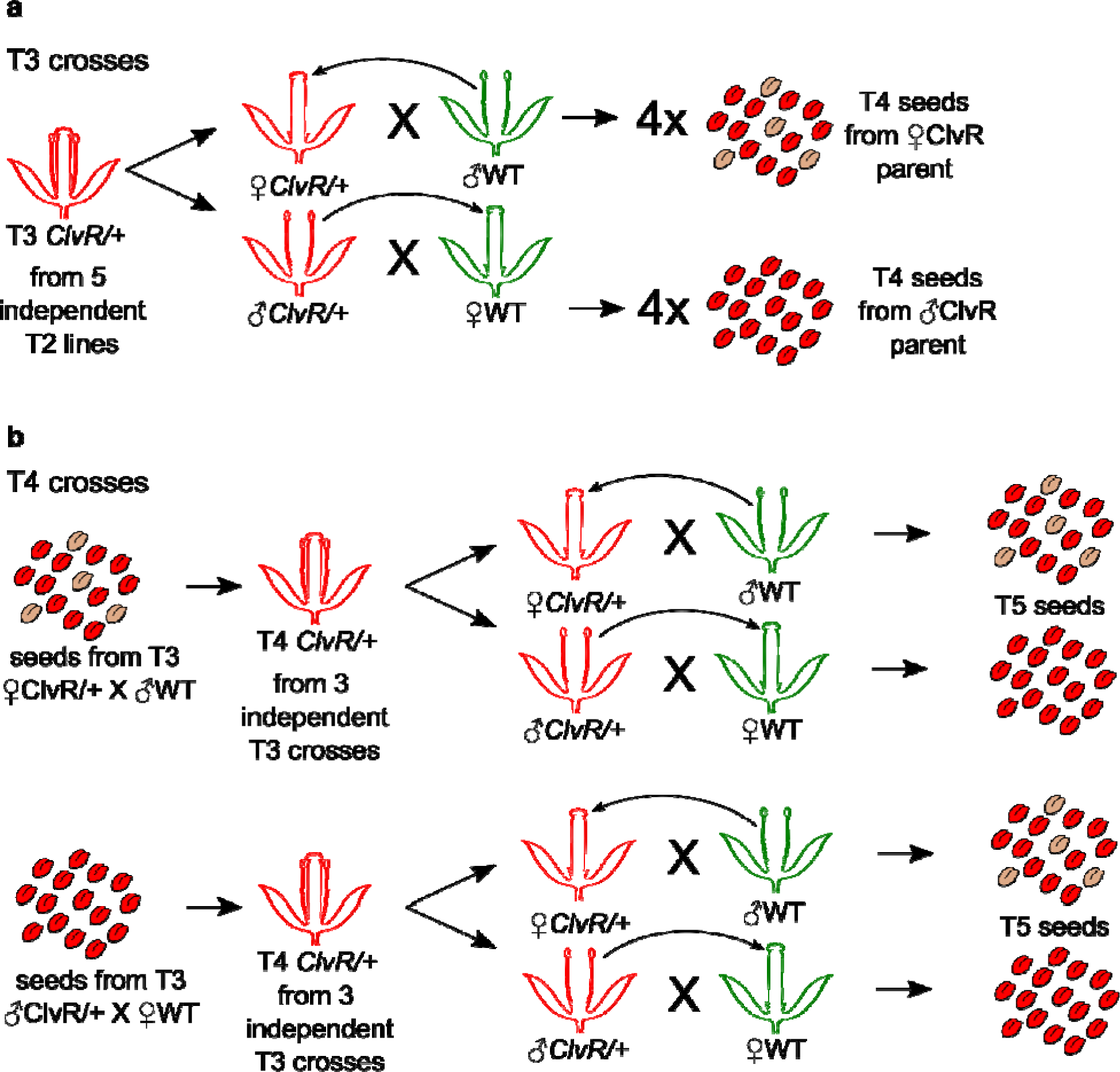
Crossing scheme for (a) T3 and (b) T4 crosses discussed in text and Fig. 2 and 3. **(a)** We selected 5 independent *ClvR^ubq^* lines that showed 100% ClvR in the T2 self cross. Pollen of T2 plants was outcrossed to WT to generate T3 heterozyogtes. For each of these 5 independent lines we set up reciprocal crosses to WT with 4 plants per line (4 crosses/siliques per plant). **(b)** For 1 of the line from (A) *ClvR*^ubq7^ we repeated the reciprocal crosses, with seeds coming from a ♀*ClvR*/+ or ♂*ClvR*/+ parent. For each of these we again crossed 4 plants (4 crosses/siliques per plant).

**Extended Data Fig. 6:**
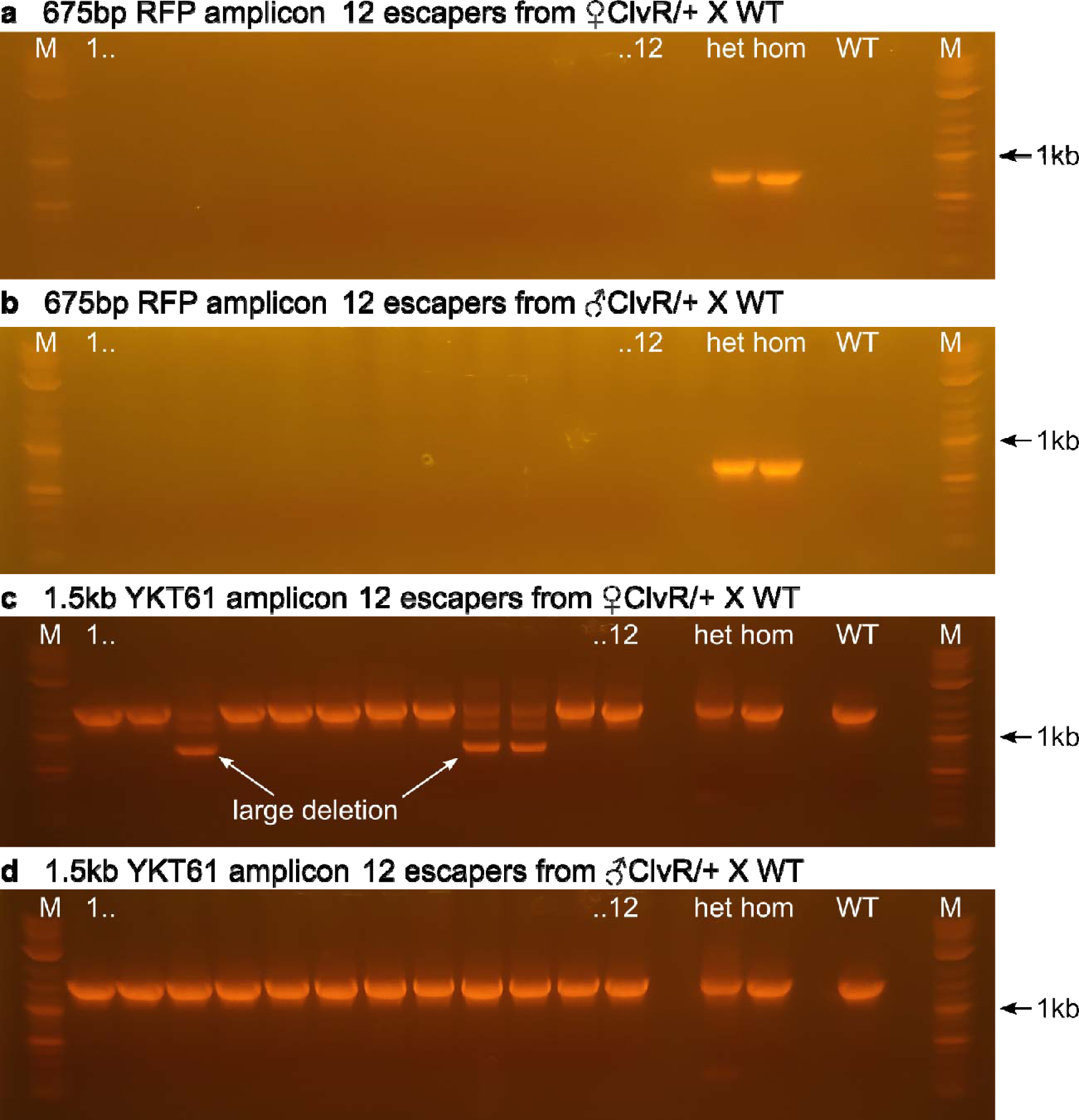
**(a-b)** PCR amplifications of a 675 bp DNA fragment of the RFP marker for escapers from a ♀*ClvR*/+ X WT **(a)** or ♂*ClvR*/+ X WT **(b)** cross. Hetero- and homozygous (het, hom) *ClvR* plants were used as positive controls, WT as negative control. Only *ClvR*-bearing plants showed the RFP band. **(c-d)** Control PCRs on the same DNA samples as in **a** and **b**, in which the YKT61 target region was amplified. Note some female escapers in **c** had larger deletions.

**Extended Data Fig. 7:**
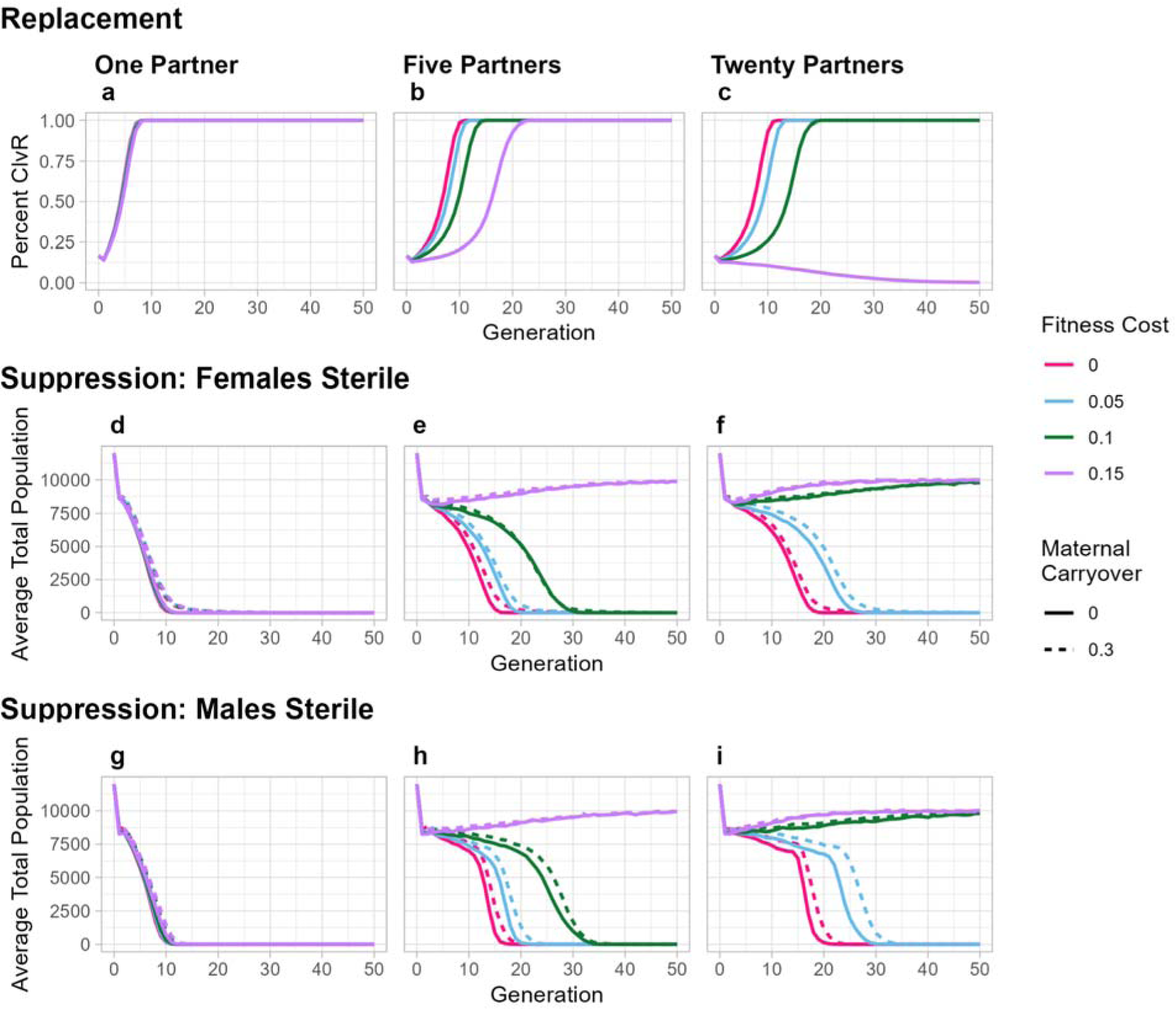
Predicted behavior of *ClvR* for population modification and suppression. **(a-c)** Population modification. *ClvR* is introduced as homozygous males at a frequency of 20% of the starting population, which is at carrying capacity, 10,000 individuals. The mating system is monogamous **(a)**, or polyandrous, with 5 males each providing 1/5th of the pollen needed to fertilize all ovules of an individual female **(b)**, or 20 males each providing 1/20th of the pollen needed **(c)**. Fitness costs are incurred by gametes (a probability of not being able to participate in fertilization, if chosen by the model). Maternal carryover is set to zero. Lines represent the average of 10 runs. **(d-f)** Population suppression with a transgene inserted into a recessive locus required for female sporophyte fertility. *ClvR* is introduced as above, at a frequency of 20%. The mating system is monogamous **(d)**, or polyandrous, with 5 males each providing 1/5th of the pollen needed to fertilize all ovules of an individual female **(e)**, or 20 males each providing 1/20th of the pollen needed **(f)**. Fitness costs are as above. Maternal carryover is set to zero or 30% (the approximate value observed in our experiments with *ClvR*^ubq^). **(g-i)**. As with **d-f**, but with the *ClvR* inserted into a locus required for male sporophyte fertility. For these simulations homozygous females were released into the population since homozygous males are sterile. Lines represent the average of 10 runs. For all panels compare with 10% introduction frequency data shown in Fig. 5.

**Extended Data Fig. 8:**
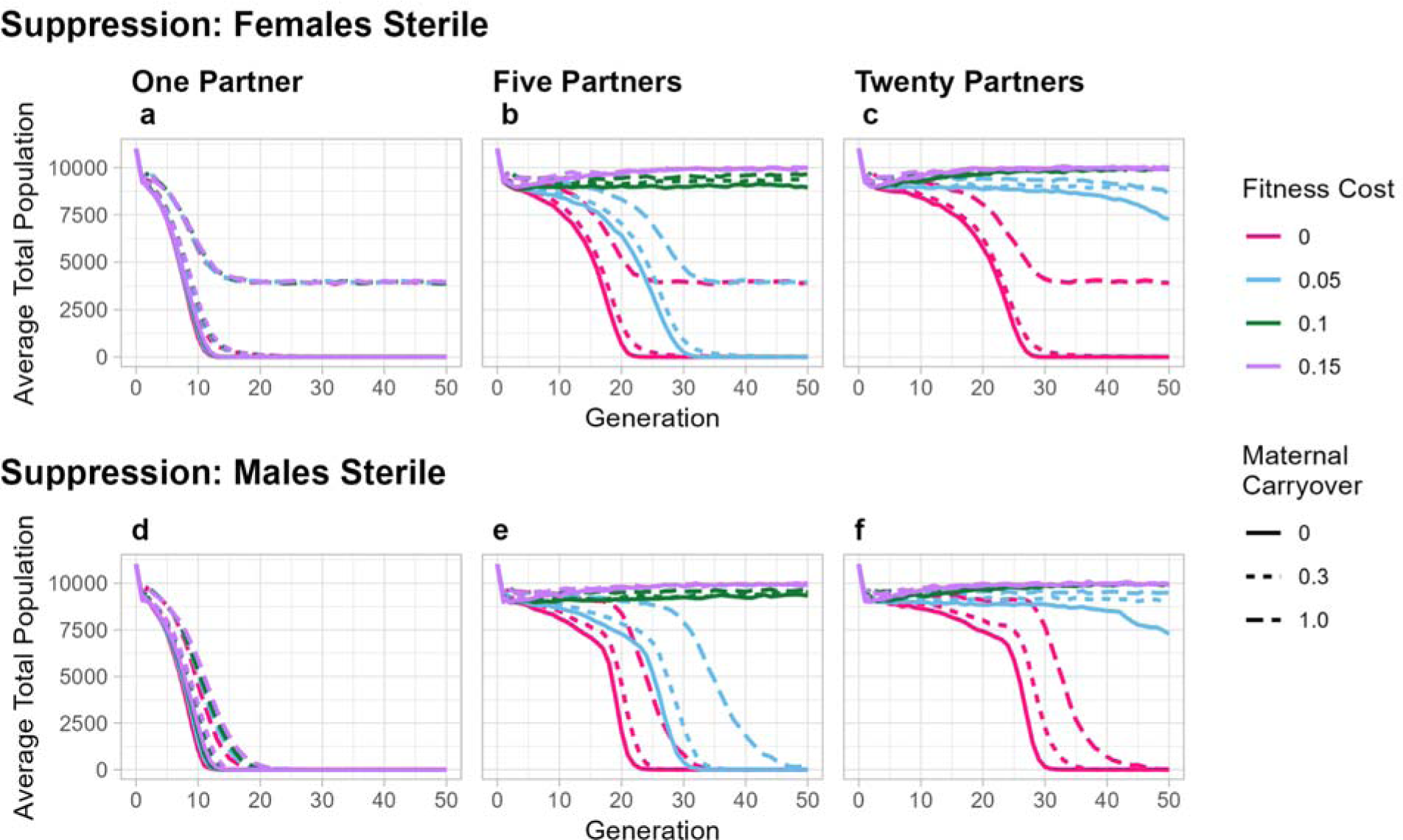
Predicted behavior of *ClvR* for population suppression with 100% maternal carryover. *ClvR* is introduced at a frequency of 10%, and is present in a female fertility locus **(a-c)** or a male fertility locus **(d-f)**, thereby creating a LOF allele. **(a-c)** When *ClvR* is located in a gene required for female sporophyte fertility high levels of maternal carryover prevent population extinction. **(d-f)** In contrast, when *ClvR* is located in a gene required for male sporophyte fertility, population extinction is slowed but not prevented.

**Extended Data Fig. 9:**
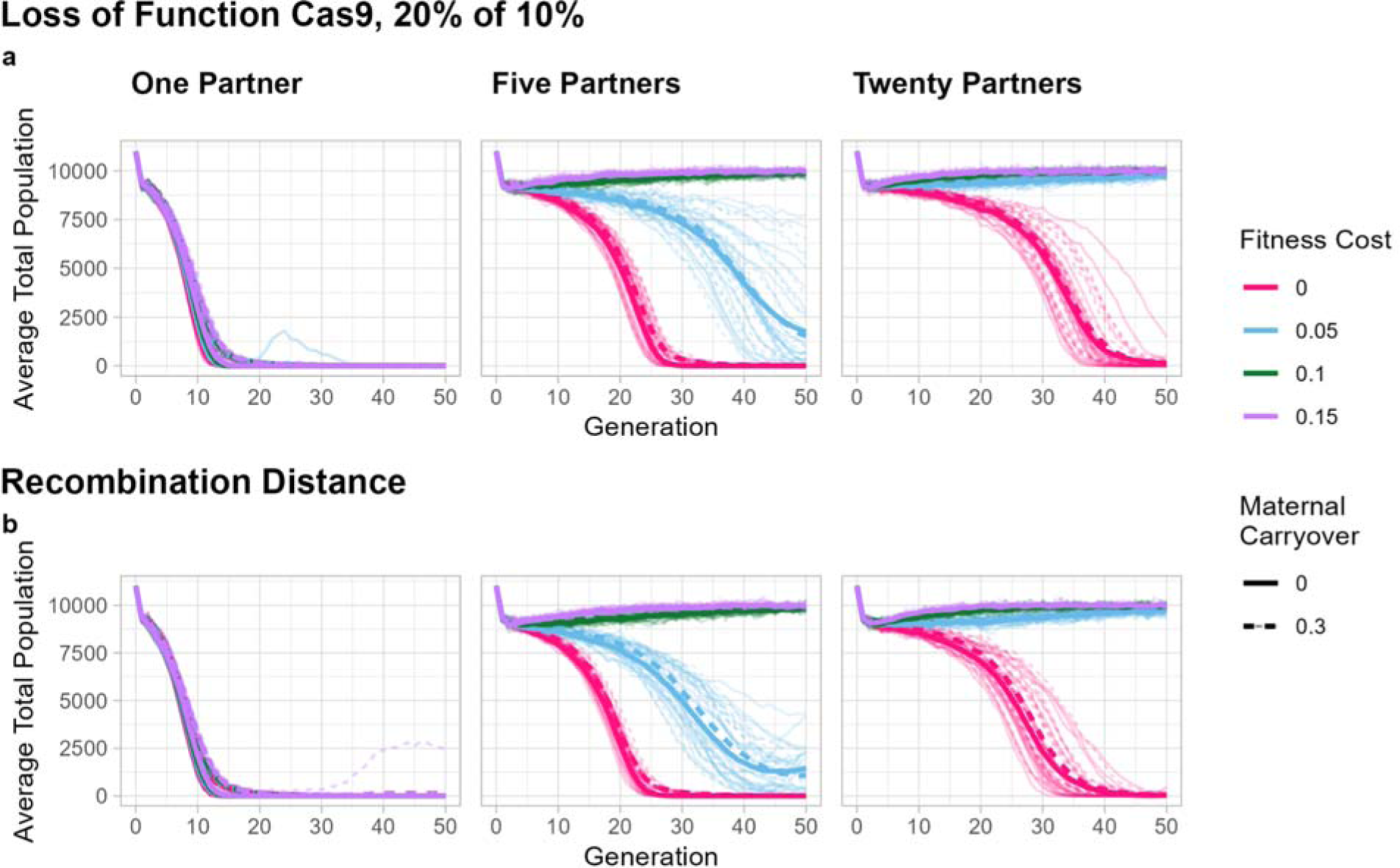
Effects of higher levels of elements lacking Cas9 on *ClvR*-mediated population suppression. **(a)** *ClvR*, located inside a gene required in the sporophyte for female fertility, is introduced at a frequency of 10%, with 20% of these elements lacking Cas9. Individual runs are shown in thin lines and the average as a thick line. **(b)** *ClvR*, located in a gene required in the sporophyte for female fertility, is introduced at a frequency of 10%. Cas9 is located 1 map unit (1% recombination rate) away from the Rescue/gRNAs. Multiple individual runs fail to go to extinction while others that do go to extinction take much longer than under the conditions shown in Fig. 5.

**Extended Data Fig. 10:**
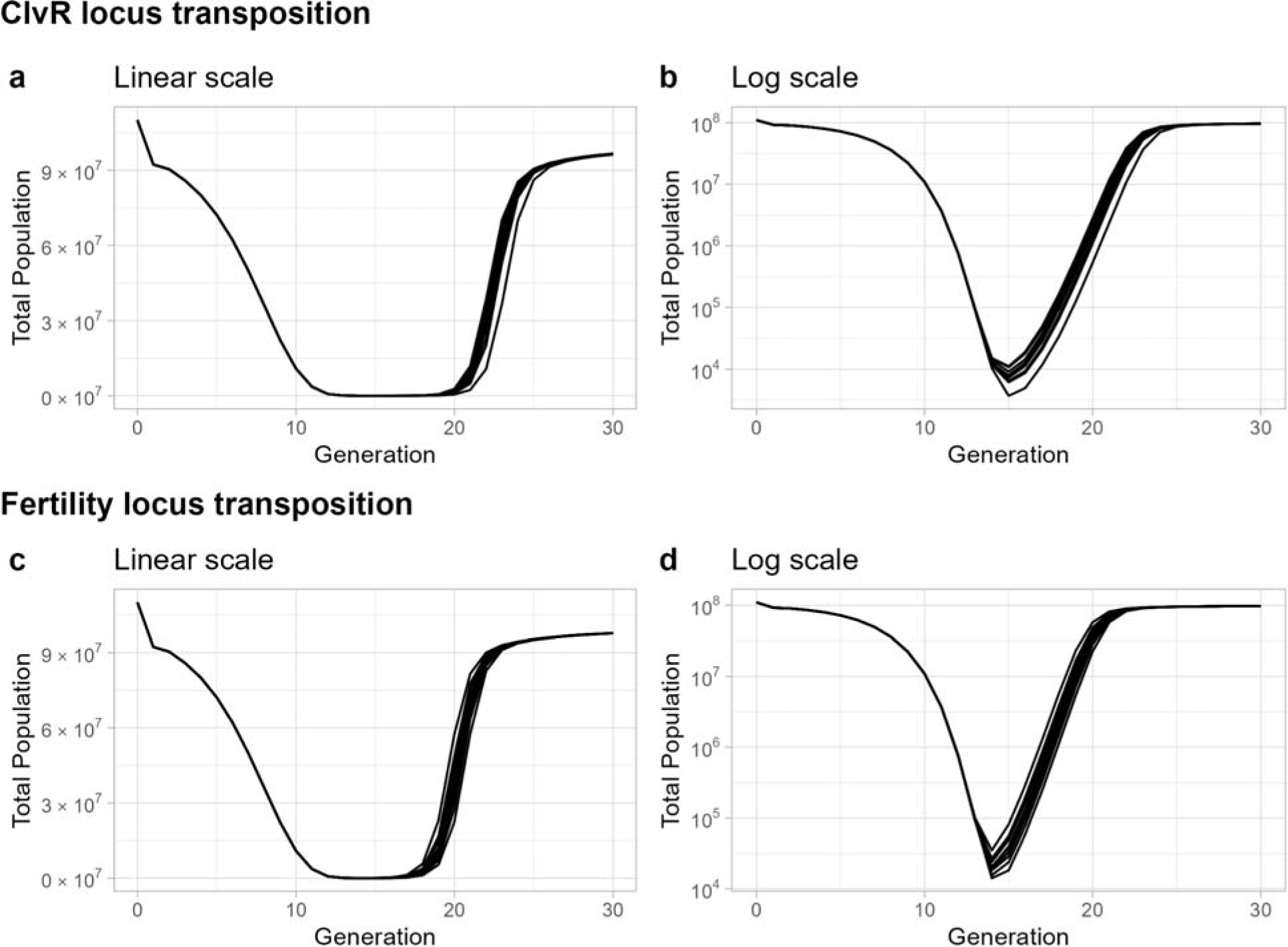
Movement of a population suppression *ClvR* or a WT allele of the sporophyte fertility gene to a new unlinked location negatively affects population suppression. **(a-b)** *ClvR*, located inside a gene required in the sporophyte for female fertility, is introduced at a frequency of 10%, with 1×10^−6^ of the *ClvR* elements having been transposed to a new unlinked locus, which is not required for female fertility. Individual runs are shown here, in a linear scale **(a)** and logarithmic scale **(b)**. **(c-d)** *ClvR*, located inside a gene required in the sporophyte for female fertility, is introduced at a frequency of 10%. Additionally, a translocated WT version of the fertility locus is present at a third locus, not associated with *ClvR*, at an allele frequency of 5×10^−7^ (100 out of 100,000,000 individuals are heterozygous for this translocated fertility gene), such that individuals with that gene may be both *ClvR* homozygous and fertile. Individuals runs are shown on a linear scale **(c)** and a logarithmic scale **(d)**.

**Extended Data Fig. 11:**
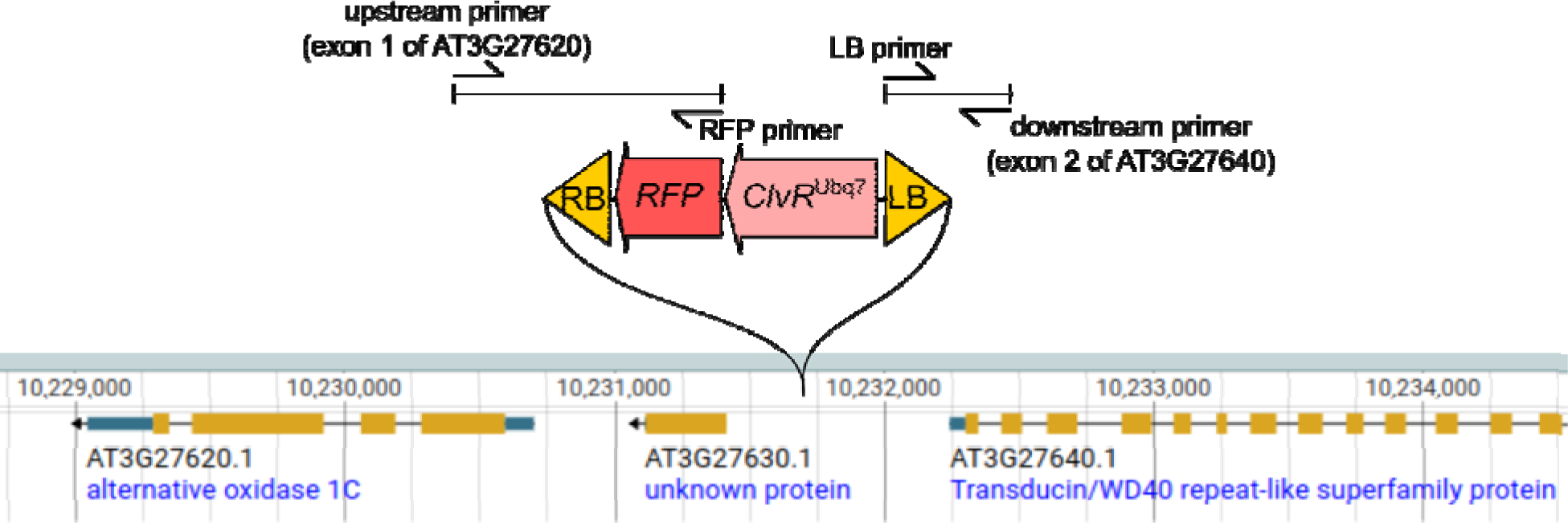
T-DNA insertion site of *ClvR*^ubq^^7^. *ClvR*^ubq7^ is inserted in the intergenic region between AT3G27630 and AT3G27640 (TAIR10, Chr3: 10231731). The upstream breakpoint was amplified in a PCR with a primer binding in exon 1 of AT3G27620 and a construct specific primer binding at the start of the RFP marker gene. The downstream breakpoint was amplified with a primer binding in exon 2 of AT3G27640 and a construct specific primer binding in the LB region of the T-DNA. Sequences of the PCR amplicons are in Supplementary Data 2.

**Extended Data Fig. 12:**
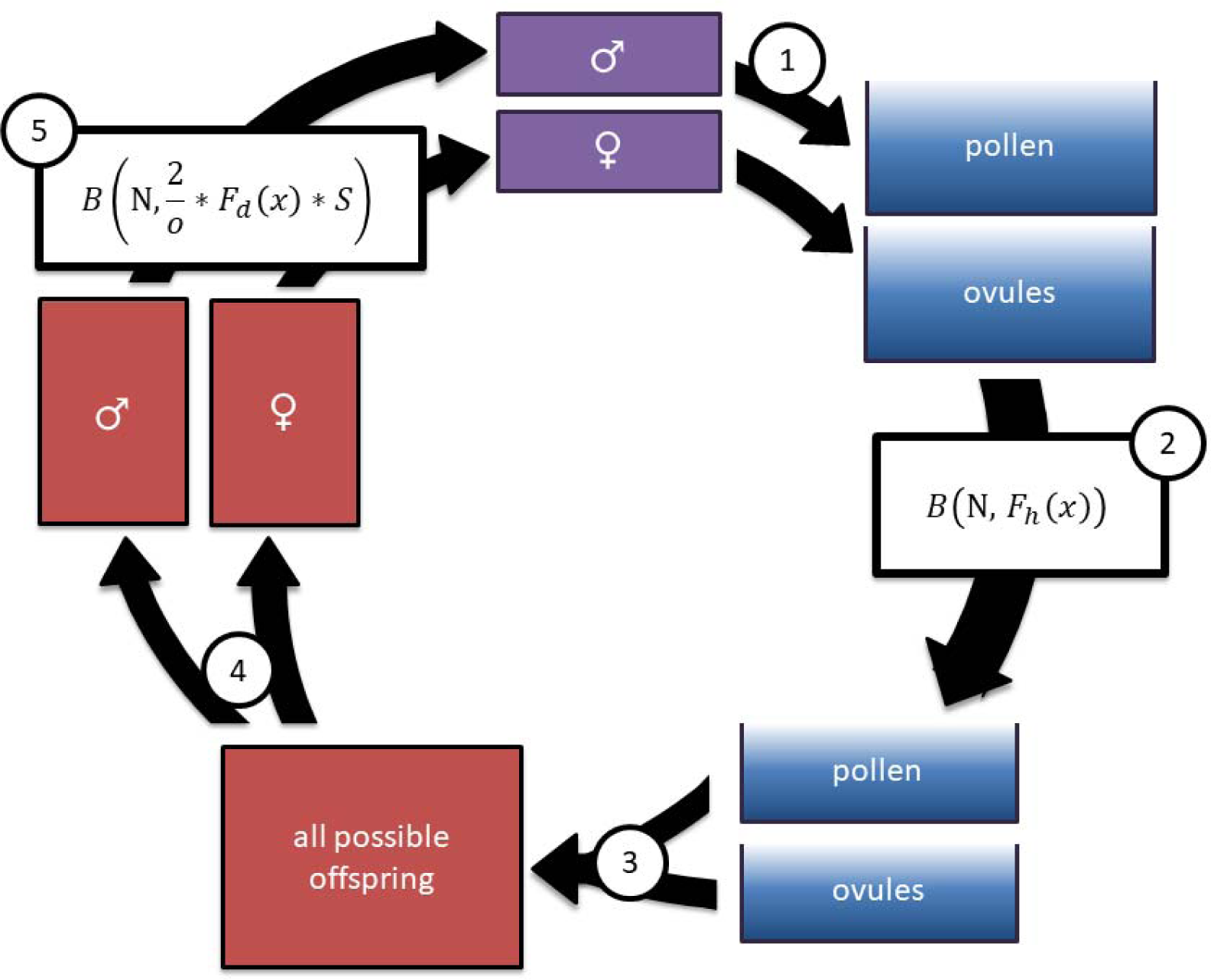
Modeling Process. Each generation starts with a pool of adult individuals, shown at the top in purple. This information is stored as a list where each index represents a possible genotype, and the associated value represents the number of individuals of that genotype. **(1)** For each mating, a single mother produces a pool of ovules and one (monogamous) or multiple (polygamous) males produce a single pool of pollen. **(2)** The number of ovules and pollen are reduced, based on their haploid fitness costs *F*_h_ such that a pollen with no fitness costs will have a 100% chance of survival, and a pollen with fitness cost 0.1 will only have a 90% chance of surviving. **(3)** From the pool of possible pollen and ovules, each ovule is mated with a single pollen to produce a possible offspring. **(4)** The possible offspring are randomly grouped as male or female **(5)**. The number of possible offspring is reduced to only those that survive. For each mating, the base chance of survival is 2 over the number of expected offspring *o*, as we expect each mating between a female and a male to produce 2 offspring when the population is at carrying capacity. This probability of survival is further modified by *F_d_*, the fitness of the individual, and *S*. *S* denotes the density dependence function *S(P) = g / [1 + (g-1)* P/K]* where *P* is the parent generation’s population size, and *K* is the carrying capacity. *S* ranges from some growth factor *g* = 6 at low densities, to 1 at densities near carrying capacity. The offspring that survive become the parents of the next generation.

**Extended Data Table 1:** Sequencing results of the YKT61 locus for various *ClvR* and escaper genotypes. “+” indicates insertions, “-” indicates deletions with numbers and type of bases deleted/inserted, WT indicates unaltered or wildtype sequences.

| Heterozygous *ClvR* | gRNA1 | gRNA2 | gRNA3 | gRNA4 |
| --- | --- | --- | --- | --- |
| Ubq7_T4_het1 | -1T/+1T | WT | +1T | WT |
| Ubq7_T4_het2 | +1T | +1T / -1G | +1T | +1C |
| Ubq7_T4_het3 | +1T | WT | +1T | +1C |
| Ubq7_T4_het4 | -95 / +1T | WT | +1T | +1A |
| Ubq7_T4_het5 | +1T / -1T | +1C | +1T / -1 T | -1C/+1A |
| inbred 1 generation | gRNA1 | gRNA2 | gRNA3 | gRNA4 |
| Ubq7_T5_hom1 | +1T / +1A | -1G | -1T | +1C |
| Ubq7_T5_hom2 | +1T | -5GTCAA | +1T | -1C |
| Ubq7_T5_hom3 | +1T / +1A | -1G | -1T | +1C |
| Ubq7_T5_hom4 | +1T | -2GC / +1A | -1T | +1C |

| Escapers from females | gRNA1 | gRNA2 | gRNA3 | gRNA4 |
| --- | --- | --- | --- | --- |
| Ubq7_T4_esc1 | WT | WT | WT | WT |
| Ubq7_T4_esc2 | WT | WT | WT | -35 |
| Ubq7_T4_esc3 | +1T | WT | WT | +1C |
| Ubq7_T4_esc4 | +1T | WT | WT | +1C |
| Ubq7_T4_esc5 | -1T | WT | WT | WT |
| Ubq7_T4_esc6 | -1T | WT | WT | WT |
| Ubq7_T5_esc1 | +51bp indel | WT | WT | +1C |
| Ubq7_T5_esc2 | +1T | WT | +1T | +1T |
| Ubq7_T5_esc3 | +1T | WT | deletion between gRNA3 and 4 |  |
| Ubq7_T5_esc4 | +1T | WT | deletion between gRNA3 and 4 |  |
| Escapers from males | gRNA1 | gRNA2 | gRNA3 | gRNA4 |
| Ubq7_T5_esc1 | WT | WT | WT | WT |
| Ubq7_T5_esc2 | WT | WT | WT | WT |
| Ubq7_T5_esc3 | WT | WT | WT | WT |
| Ubq7_T5_esc4 | WT | WT | WT | WT |
| Ubq7_T5_esc5 | WT | WT | WT | WT |
| Ubq7_T5_esc6 | WT | WT | WT | WT |

**Extended Data Table 2:** Self crosses of escapers ♀*ClvR*/+ X ♂WT cross.

| T5 escapers from female *ClvR/+* |  |  |  |
| --- | --- | --- | --- |
| escaper Plant | Seeds | aborted | % survival |
| Ubq7_esc1A | 13 | 21 | 38.2 |
| Ubq7_esc1B | 16 | 16 | 50.0 |
| Ubq7_esc1C | 12 | 14 | 46.2 |
| Ubq7_esc2A | 19 | 19 | 50.0 |
| Ubq7_esc2B | 23 | 25 | 47.9 |
| Ubq7_esc2C | 17 | 20 | 45.9 |
| Ubq7_esc3A | 23 | 25 | 47.9 |
| Ubq7_esc3B | 27 | 20 | 57.4 |
| Ubq7_esc3C | 24 | 21 | 53.3 |
| Ubq7_esc4A | 23 | 21 | 52.3 |
| Ubq7_esc4B | 22 | 19 | 53.7 |
| Ubq7_esc4C | 25 | 21 | 54.3 |
| Ubq7_esc5A | 24 | 20 | 54.5 |
| Ubq7_esc5B | 22 | 26 | 45.8 |
| Ubq7_esc5C | 26 | 18 | 59.1 |
| Ubq7_esc6A | 12 | 24 | 33.3 |
| Ubq7_esc6B | 20 | 22 | 47.6 |
| Ubq7_esc6C | 18 | 21 | 46.2 |
| **Total** | **366** | **373** | **49.1** |

T5 escapers from male *ClvR/+*
| escaper Plant | Seeds | aborted | % survival |
| --- | --- | --- | --- |
| Ubq7_esc1A | 51 | 0 | 100.0 |
| Ubq7_esc1B | 53 | 1 | 98.1 |
| Ubq7_esc1C | 49 | 1 | 98.0 |
| Ubq7_esc2A | 52 | 0 | 100.0 |
| Ubq7_esc2B | 50 | 1 | 98.0 |
| Ubq7_esc2C | 50 | 0 | 100.0 |
| Ubq7_esc3A | 40 | 1 | 97.6 |
| Ubq7_esc3B | 44 | 2 | 95.7 |
| Ubq7_esc3C | 41 | 1 | 97.6 |
| **Total** | **430** | **7** | **98.8** |

control, wildtype self cross: WT X WT
| Plant | Seeds | aborted | % survival |
| --- | --- | --- | --- |
| WT1A | 48 | 0 | 100.0 |
| WT1B | 47 | 1 | 97.9 |
| WT1C | 52 | 1 | 98.1 |
| WT2A | 46 | 0 | 100.0 |
| WT2B | 44 | 0 | 100.0 |
| WT2C | 40 | 1 | 97.6 |
| WT3A | 50 | 1 | 98.0 |
| WT3B | 47 | 1 | 97.9 |
| WT3C | 50 | 0 | 100.0 |
| WT4A | 46 | 0 | 100.0 |
| WT4B | 51 | 0 | 100.0 |
| WT4C | 48 | 1 | 98.0 |
| WT5A | 47 | 1 | 97.9 |
| WT5B | 44 | 1 | 97.8 |
| WT5C | 45 | 1 | 97.8 |
| **Total** | **705** | **9** | **98.7** |

homozygous *ClvR* self cross: *ClvR/ClvR* X *ClvR/ClvR*
| Plant | Seeds | aborted | % survival |
| --- | --- | --- | --- |
| Ubq7_hom1A7 | 52 | 0 | 100.0 |
| Ubq7_hom1B | 50 | 1 | 98.0 |
| Ubq7_hom1C | 47 | 1 | 97.9 |
| Ubq7_hom2A | 48 | 0 | 100.0 |
| Ubq7_hom2B | 54 | 1 | 98.2 |
| Ubq7_hom2C | 49 | 1 | 98.0 |
| Ubq7_hom3A | 41 | 2 | 95.3 |
| Ubq7_hom3B | 40 | 2 | 95.2 |
| Ubq7_hom3C | 40 | 1 | 97.6 |
| **Total** | **421** | **9** | **97.9** |

